# Imbalanced gut microbiota predicts and drives the progression of nonalcoholic fatty liver disease and nonalcoholic steatohepatitis in a fast-food diet mouse model

**DOI:** 10.1101/2023.01.09.523249

**Authors:** Na Fei, Sawako Miyoshi, Jake B. Hermanson, Jun Miyoshi, Bingqing Xie, Orlando DeLeon, Maximilian Hawkins, William Charlton, Mark D’Souza, John Hart, Dinanath Sulakhe, Kristina B. Martinez-Guryn, Eugene B. Chang, Michael R. Charlton, Vanessa A. Leone

## Abstract

Nonalcoholic fatty liver disease (NAFLD) is multifactorial in nature, affecting over a billion people worldwide. The gut microbiome has emerged as an associative factor in NAFLD, yet mechanistic contributions are unclear. Here, we show fast food (FF) diets containing high fat, added cholesterol, and fructose/glucose drinking water differentially impact short- vs. long-term NAFLD severity and progression in conventionally-raised, but not germ-free mice. Correlation and machine learning analyses independently demonstrate FF diets induce early and specific gut microbiota changes that are predictive of NAFLD indicators, with corresponding microbial community instability relative to control-fed mice. Shotgun metagenomics showed FF diets containing high cholesterol elevate fecal pro-inflammatory effectors over time, relating to a reshaping of host hepatic metabolic and inflammatory transcriptomes. FF diet-induced gut dysbiosis precedes onset and is highly predictive of NAFLD outcomes, providing potential insights into microbially-based pathogenesis and therapeutics.

**Highlights:**

- Germ-free mice are protected from fast-food diet-induced NAFLD.
- Fast-food diets rapidly shift gut microbiota composition and function.
- Increasing dietary cholesterol exacerbates hepatic inflammation only in SPF mice.
- Fast-food diet-induced gut dysbiosis precedes and predicts late-stage NAFLD severity.

## Introduction

Nonalcoholic fatty liver disease (NAFLD), and its more severe form, nonalcoholic steatohepatitis (NASH), are the most common chronic liver diseases worldwide and have become the leading indications of liver transplantation in certain populations in the United States (Ballestri et al. 2015, Li et al. 2019). Similar to common comorbidities of obesity and diabetes, the emerging global epidemic of NAFLD and NASH has placed a major strain on healthcare systems worldwide (Estes et al. 2018); mechanisms remain unclear, and while several interventions are in clinical trials, current therapeutic options for NAFLD and NASH management are limited with no approved treatments (Benedict and Zhang 2017). Finding a means to prevent or reverse NAFLD and NASH would have a profound clinical impact, improving the patient prognosis and reducing the overall healthcare burden.

Over the past decade, a rapidly growing body of evidence indicates the gut–liver axis plays a crucial role in NAFLD and NASH pathophysiology (Roychowdhury, Selvakumar and Cresci 2018, Sharpton et al. 2021). Both preclinical and human clinical studies suggest gut microbes significantly contribute to development of metabolic diseases, including obesity, diabetes, and NAFLD (Lynch, Chan and Drake 2017, Roychowdhury, Selvakumar and Cresci 2018), demonstrating overlapping microbial signatures that can differentiate between NAFLD and NASH subjects vs. healthy controls (Kalhan et al. 2011, Loomba et al. 2017, Caussy et al. 2019). NAFLD is frequently associated with increased abundance of the phylum Proteobacteria, mainly implicating genera and species in the family *Enterobacteriaceae* (Fei et al. 2020, Hoyles et al. 2018, Loomba et al. 2017, Raman et al. 2013, Shen et al. 2017), along with decreased abundance of the phylum *Firmicutes* and families *Rikenellaceae* and *Rumminoccaceae* (Del Chierico et al. 2017, Raman et al. 2013, Shen et al. 2017, Zhu et al. 2013).

While human studies remain associative, animal models using fecal microbiota transplantation (FMT) with full microbial communities from NAFLD vs. healthy subjects or mono-association with a single gut bacteria demonstrate a causal role of gut microbiota in disease development (Fei et al., 2020, Zhang et al. 2021, Natividad et al. 2018). For instance, FMT from NAFLD and NASH conventionally-raised mice into germ-free (GF) mice completely devoid of all microbiota reproduced some NAFLD histological features, including steatosis and an increase in hepatic triglyceride content regardless of dietary intake. Further, FMT from mice with NAFLD, but not from healthy controls, augmented hepatic gene expression in recipient animals (Henao-Mejia et al. 2012, Le Roy et al. 2013). Additionally, FMT from NAFLD/NASH patients into GF mice triggered similar NAFLD/NASH phenotypes, including elevated circulating lipopolysaccharide (LPS) concentration under both undefine, grain-based chow (Hoyles et al. 2018) and high fat (HF) diet feeding (Chiu et al. 2017). Mono-association of GF mice with gut bacteria strains isolated from human NAFLD/NASH patients, such as *Enterobacter cloacae* B29, *E. coli* PY102, or *Klebsiella pneumonia* A7 can induce NAFLD upon exposure to high-fat feeding (Fei et al. 2020), indicating a complex interaction between the host, gut microbes, and diet. Mechanistically, a common feature of these studies using FMT or mono-association is elevated circulating microbially-derived Lipopolysaccharide (LPS) levels that may further contribute to NAFLD development and progression to NASH via enhanced inflammation and fibrosis. Elevated circulating LPS is a key feature of increased intestinal permeability, which can trigger tissue and systemic inflammation (Cani et al. 2007, Carpino et al. 2020, Mao et al. 2015, Wang et al. 2021). Collectively, this evidence indicates an essential role of gut microbiota in NAFLD and NASH development. However, despite these findings, contributions of key dietary components to the onset of gut dysbiosis in the context of gut-liver axis disruption and NAFLD development remains limited.

NAFLD and NASH rodent models provide increasing insight into gut dysbiosis as a causative factor in disease; however, the utility and translational aspects are limited, including an inability to faithfully recapitulate human dietary intake. For instance, choline-deficient high-fat diet is widely used as a NASH model (Rinella et al. 2008), yet this does not correspond to human dietary intake. Dietary cholesterol is an established contributor to NAFLD in humans (Noureddin et al. 2000, Mokhtari et al. 2017, Yasutake et al. 2009, Loannou et al. 2009); however, few murine studies have considered this factor in diet design. Previous studies show that the murine fast-food model (FF, high fat, added cholesterol diet plus fructose/glucose water) recapitulates human dietary intake (35.8% kcal from total fat [CDC in the 2015-2018 survey], 290 mg/day of cholesterol [NHANES in the 2013–2014 survey]) with corresponding features of human NAFLD/NASH development, progression, and severity as well as metabolic facets of NAFLD/NASH, such as obesity, insulin resistance and dyslipidemia (Charlton et al. 2011, Krishnan et al. 2014). These studies revealed that mice fed FF diet developed fibrotic NASH, while mice fed HF plus sugar water with no added cholesterol only developed obesity and simple liver steatosis without fibrosis (Charlton et al. 2011, Krishnan et al. 2014). Two factors remain unexplored – 1) the role of gut microbes in FF diet-induced NAFLD/NASH development and 2) their interaction with the essential dietary component, cholesterol. Here, the central focus of the current study was to disentangle these complex host-microbe-diet interactions and gain insights into their timing in NAFLD development and the transition to NASH. We hypothesized that FF diets containing increasing levels of dietary cholesterol would induce an earlier onset of gut dysbiosis, which contributes to elevated biomarkers of NAFLD and NASH and, ultimately, worse disease outcomes.

Using a combination of conventionally-raised, specific-pathogen-free (SPF) and GF mice fed FF diets, we reveal that cholesterol level in the FF diet promotes a rapid reshaping of the gut microbiota that precedes disease onset and contributes to a faster progression of NAFLD to NASH. This early diet-induced gut dysbiosis appears to be highly predictive of late-stage disease, which could provide insights into the timing of possible therapeutic interventions to reverse disease course.

## Results

### Fast food (FF) dietary cholesterol level differentially impacts short- and long-term disease severity and progression of NAFLD/NASH in SPF, but not GF, mice

First, we fed low and high-fat purified diets containing two levels of added dietary cholesterol to SPF and GF mice to test whether FF-diet-induced gut dysbiosis plays an essential role in the development and progression of NAFLD (**Figure 1A**). Over the 24-week observation period, SPF mice fed HF or FF diet, regardless of cholesterol level, exhibited significantly increased percent body weight gain relative to low-fat (LF)-fed SPF counterparts, while no appreciable differences in weight gain were observed over time or between diet groups in GF conditions (**Figure 1B**). The increased weight gain in HF and FF-fed SPF vs. GF animals was not explained by differences in caloric intake, as both groups, regardless of microbial status, ate and drank nearly the same amount of kcals across all diets throughout the experiment (**Figure S1A, B**). These results confirmed gut microbes are required for HF and FF-diet-induced weight gain regardless of cholesterol level that is not explained by differences in caloric consumption.

**Figure 1.**
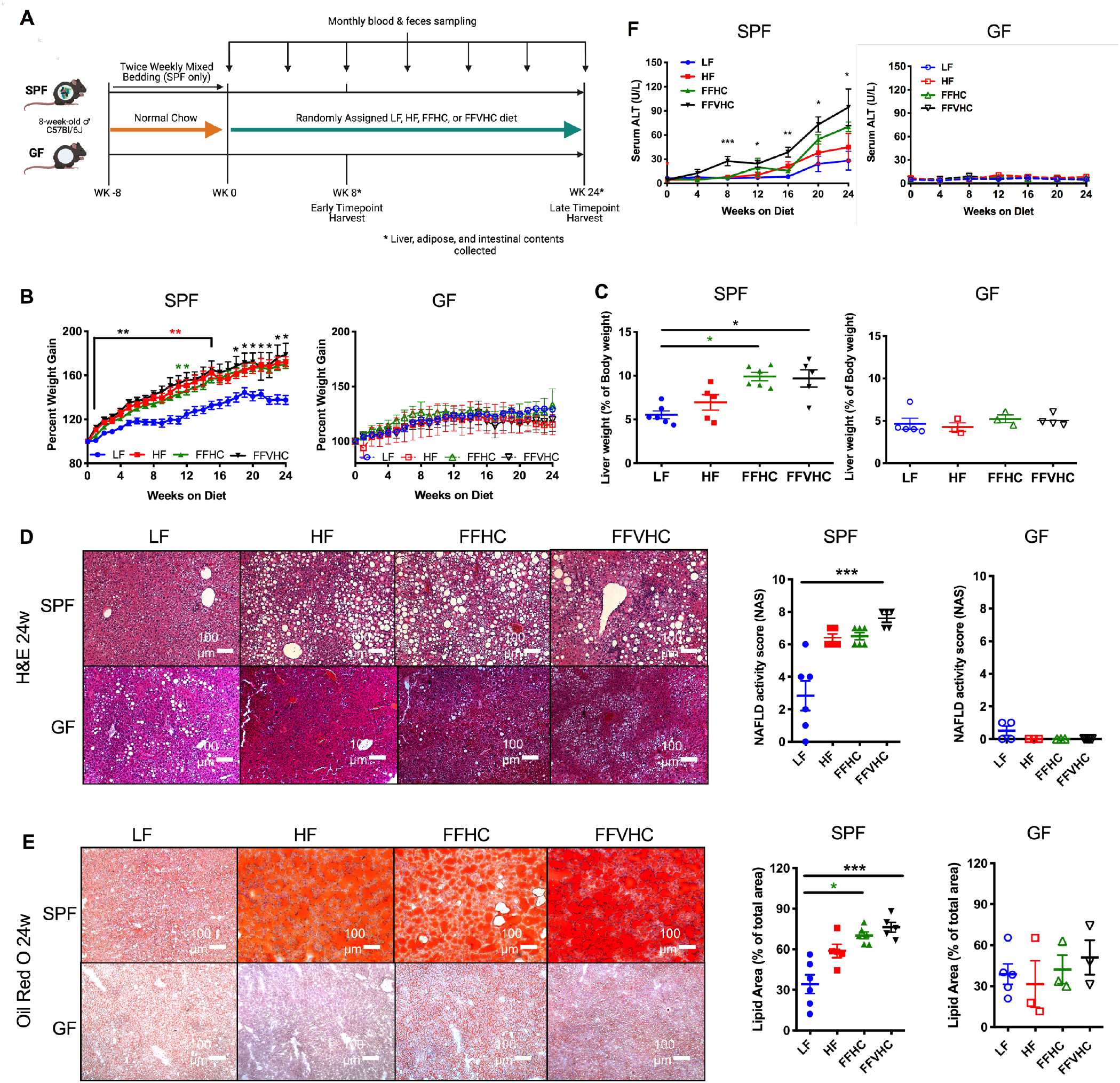
Dietary cholesterol level differentially impacts progression of NAFLD/NASH in specific-pathogen-free (SPF) but not germ-free (GF) mice after 24 weeks of feeding. (**A**) Fast-food diet mouse model depicting experimental timeline indicating key timepoints and tissue collections in SPF and GF mice. (**B**) Percent body weight gain and (**C**) Liver weight expressed as % of body weight at 24 weeks in SPF and GF mice. (**D**) Formalin-fixed liver hematoxylin and eosin stain (H&E, right) and corresponding graphs represent NAFLD activity scores (NAS, left) of SPF and GF mice. All microscopy images were at 100x magnification. (**E**) Oil Red-O staining (left) and corresponding graph representing lipid area quantification in SPF and GF mice at 24 weeks. Representative images for all histology are shown for livers from each group. All microscopy images were at 100x magnification, Scale bars = 100 mm. (**F**) Time course of alanine aminotransferase (ALT) levels in peripheral plasma of SPF and GF mice. Data represent means ± S.E.M. *P < 0.05, **P < 0.01, ***P < 0.001 via nonparametric Kruskal-Wallis one-way ANOVA test followed by Tukey’s multiple comparisons between groups at each week or time point. n=4-6 per group. Star color represents the treatment group exhibiting significant differences.

After 8 and 24 weeks during the observation period, we next assessed how FF diet, as well as the presence or absence of gut microbiota, influenced NAFLD/NASH disease outcomes. Expressed as a percent of body weight, no significant differences in adipose tissue deposition (retroperitoneal, gonadal, inguinal, and mesenteric fat) were detected in SPF or GF mice across diets at 24 weeks (**Figure S1C**). Gross assessment of livers revealed that FF diets, regardless of cholesterol level, significantly increased liver weight (expressed as % of body weight) in SPF FFHC and FFVHC mice relative to LF-fed counterparts at 24 weeks (**Figure 1C**), although trends were evident for FFHC and FFVHC-fed mice as early as 8 weeks after onset of diet (**Figure S1D**). No significant differences at either time point were observed in liver weight between diet groups in GF conditions (**Figure 1C, Figure S1D**). Blinded histological evaluation of H&E-stained formalin-fixed liver sections by a trained pathologist revealed HF-, FFHC-, and FFVHC-fed SPF mice exhibited increased evidence of steatosis and inflammation relative to LF-fed SPF mice, while FFVHC induced a statistically elevated NAFLD activity score (NAS) at both 8 and 24 weeks (**Figure S1E, Figure 1D**, p<0.05 and p<0.001 - 8w: LF, 0.75; HF, 5.6; FFHC, 5.4; FFVHC, 7.8; 24w: LF, 2.8; HF, 6.4; FFHC, 6.5; FFVHC, 7.6, respectively). No significant differences in NAS were observed between any group in GF mice at either time point (**Figure S1E, Figure 1D**). Liver lipid deposition (calculated as percent lipid area relative to the total area), another hallmark feature of NAFLD and NASH determined via Oil red-O staining, was dramatically and significantly elevated in FFHC- and FFVHC-fed SPF mice relative to LF-fed SPF mice at 24 weeks (**Figure 1E**; p<0.05 and p<0.001, respectively), with evidence of significantly increased lipid deposition at 8 weeks in FFVHC-fed SPF animals (**Figure S1F**). No significant differences in lipid deposition were detected in GF mice across diets or time points (**Figure 1E, Figure S1F**).

Plasma alanine aminotransferase (ALT) level, a classic clinical indicator of liver dysfunction, was significantly elevated in FFVHC-fed SPF mice relative to LF-fed counterparts, beginning at 8 weeks of feeding, which persisted throughout the 24-week observation period (**Figure 1F, Figure S1G**). This appeared to depend on the presence of microbes, as no appreciable elevation in ALT levels was observed in GF mice regardless of diet at either time point (**Figure 1F, Figure S1G**). Taken together, these results indicate gut microbes are required for diet-induced NAFLD and NASH development, i.e., GF mice are protected against disease, and that dietary cholesterol level in conjunction with exposure to high fat and high sugar accelerates disease progression in the presence of a complex microbial community.

### Increased cholesterol level in fast food diets exacerbates gut microbiota community membership imbalances that precede NAFLD onset

We next tested whether increasing cholesterol level in the FF diet contributes to shifts in microbial community membership that correspond with NAFLD onset, i.e., during the initiation phase, progression, and severity via 16S rRNA gene amplicon sequencing. We initially assessed starting microbial communities in stool collected at week 0 (baseline), a timepoint at which all mice were fed a grain-based undefined chow before random assignment to their respective semi-purified dietary treatment and monthly throughout the experiment after diet switch. β-diversity principle component analysis (PCoA) of weighted UniFrac distances revealed no significant differences in community membership between animals at baseline, confirming successful implementation of the 8-week mixed bedding normalization procedure prior to experimental initiation (**Figure S2A**) (Miyoshi et al. 2018). As expected, high sugar drinking water coupled with a semi-purified diet, regardless of fat or cholesterol level, rapidly shifted gut microbiota community membership in a similar manner after only 4 weeks of feeding, which persisted throughout the experiment (**Figure S2A**). Mice fed LF diet tended to separate more distinctly from their HF, FFHC, and FFVHC-fed counterparts throughout the 24-week period. These data confirmed a consistent community membership at baseline that rapidly shifted in response to semi-purified diet exposure.

To investigate whether shifts in gut microbiota community membership preceded disease onset and corresponded with later-stage disease outcomes, we analyzed cecal 16S rRNA gene amplicon sequencing data after only 8 weeks of diet exposure in a subset of animals as well as after 24 weeks of feeding at the end of the observation period in the remaining animals. Weighted UniFrac distances (β-diversity) of week 24 data showed HF, FFHC, and FFVHC diets shifted cecal microbiota community membership and relative abundances of specific community members, suggesting HF diets, regardless of added cholesterol level, promote a unique community membership relative to LF-fed mice in later stages of disease (P<0.005 via ADONIS, **Figure 2A**). Similar patterns were also apparent at week 8 (P<0.001, **Figure S2B**), an early time point prior to overt disease onset.

**Figure 2.**
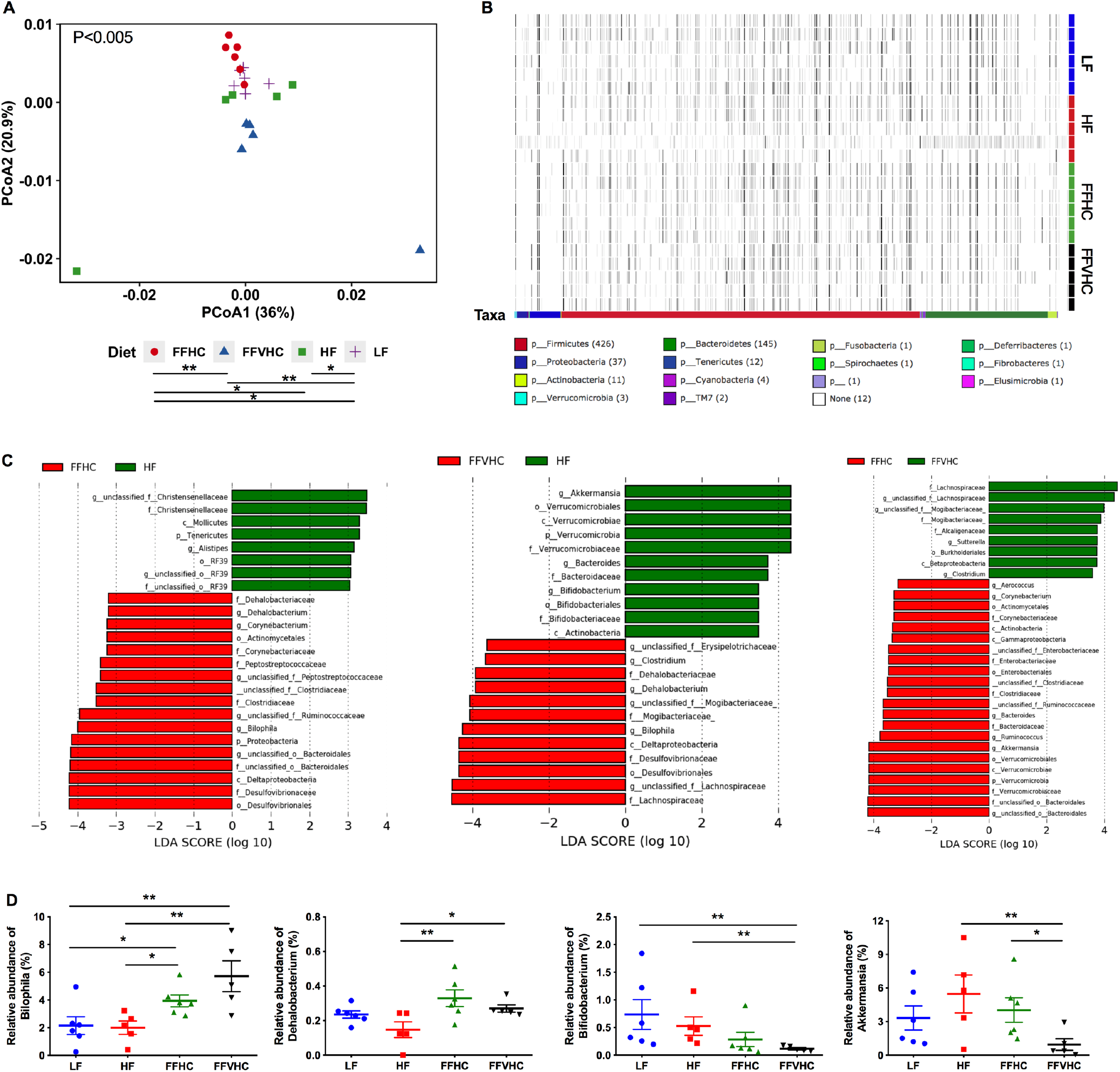
Fast-food diets significantly promote gut dysbiosis in cecal contents which is accompanied by NAFLD and NASH onset after 24 weeks of feeding. (**A**) Beta diversity Principal Coordinate Analysis (PCoA) plots for Weighted Unifrac distances at week 24 (Axis 1 = principal coordinate (PC) 1; Axis 2 = PC 2). Symbols represent individual samples. (**B**) Heat map indicating relative abundances of Amplicon Sequence Variants (ASVs) in cecal bacteria composition induced by diet. Each column in the dendrogram represents an individual ASV. Each row represents an individual sample, organized by dietary treatment to the right of the heat map. Phyla-level taxonomic assignments are indicated by the colored bars at the bottom of the heat map. (**C**) Linear discriminant analysis effect size (LEfSe) analysis (LDA score [log10] bar graphs of differentially abundant taxa from cecum of HF vs. FFHC, HF vs. FFVHC, and FFHC vs. FFVHC mice at 24 weeks. Statistically significant taxa are presented (p<0.05) with an LDA score greater than ±3. Prefixes represent taxonomic rank, i.e., phylum (p), class (c), etc. (**D**) Relative abundance of significantly changed genera induced by FF diets identified by LEfSe with LDA scores greater than ±3.0. n=4-6 per group. Data represents mean ± S.E.M. *P < 0.05, **P < 0.01, ***P < 0.001 via nonparametric Kruskal-Wallis test followed by Turkey’s multiple comparisons across all groups.

We next used Permutational multivariate analysis of variance (PERMANOVA) to perform pairwise statistical analyses of gut microbiota between diet groups, which revealed that both level of dietary fat and cholesterol resulted in significantly unique bacteria community membership (LF vs. HF [p<0.05]; LF vs. FFHC [p<0.05]; LF vs. FFVHC [p<0.01]; HF vs. FFHC [p<0.05]; FFHC vs. FFVHC [p<0.01]; **Figure 2A**). This was further demonstrated via analyses of Amplicon Sequence Variant (ASV) relative abundances via anvi’o, which showed distinct shifts in bacteria composition at the phylum level across LF, HF, and FF diets, which were further exacerbated by increasing dietary cholesterol (**Figure 2B**, heat map). Both PCoA and anvi’o analyses also showed modest but apparent differences in ASV relative abundances across diet groups after only 8 weeks following the switch to semi-purified diets (**Figure S2B, C**).

To determine ASVs that significantly contribute to differences in community membership between diet groups, we applied linear discriminant analysis effect size (LEfSe) pairwise analyses to cecal ASVs (i.e., LF vs. HF, HF vs. FFHC, FFHC vs. FFVHC, etc.). Here, we identified features across taxonomic ranks that were differentially abundant at both 8 and 24 weeks with a linear discriminant analysis (LDA) score greater than or equal to ±3.0 (**Figure 2C, Figure S2D, E, and Figure S3B**). HF diet, containing only 0.05% cholesterol, significantly enriched taxa belonging to the families *Christensenellaceae* (unclassified genus), Rikenellaceae (genus *Alistipes*), Verrucomicrobiaceae (genus *Akkermansia*), and Bifidobacteriaceae (genus Bifidobacterium) relative to FFHC and FFVHC diets, respectively, at 24 weeks. Conversely, while FF diets containing increased cholesterol level enriched for several taxa relative to HF diet at 24 weeks, the majority with a LDA score greater than or equal to ±4.0 belonged to the families Desulfovibrionaceae (genus *Bilophila*), Peptostreptococcaceae (unclassified genus), Dehalobacteriaceae (genus *Dehalobacterium*), and Lachnospiraceae (unclassified genus) (**Figure 2C**). The relative abundances of several significantly changed taxa induced by FF diets containing increasing levels of cholesterol relative to HF diet alone identified via LEfSe analysis at week 24 are shown in **Figure 2D**. Interestingly, both FFHC and FFVHC diets significantly increased *Bilophila* relative abundance in cecal contents as compared to both LF and HF-fed counterparts (P<0.05 and P<0.01, respectively), while the relative abundance of *Dehalobacterium* was only significantly elevated relative to HF, but not LF-fed mice (P<0.01 [HF vs. FFHC] and P<0.05 [HF vs. FFVHC]; **Figure 2D**, upper panels). Conversely, FFVHC diet significantly reduced *Bifidobacterium* relative abundance as compared to both LF and HF diets (P<0.01), while *Akkermansia* was significantly decreased by FFVHC as compared to HF and FFHC diets (P<0.01 and P<0.05, respectively; **Figure 2D**, bottom panels). LEfSe outcomes for comparisons between LF vs. HF, FFHC, and FFVHC are shown in **Figure S3A**, which reveals similar, yet more striking results for taxa selected by diets containing HF alone or in combination with added cholesterol.

To identify whether the observed gut microbiota changes at 24 weeks to FF diet with added cholesterol were also evident before overt disease, we performed LEfSe analysis on 8-week cecal contents (**Figure S2D, E and Figure S3B**). Here, HF diet enriched taxa belonging to the order Clostridiales (class Clostridia), genus Desulfovibrio, families Mogibacteriaceae (genus unclassified), Ruminoccocaceae (genus *Ruminococcus*), Bifidobacteriaceae (genus *Bifidobacterium*), order RF39 (unclassified genus), and class Mollicutes relative to FFHC or FFVHC diets. Conversely, FF diets containing increasing cholesterol levels enriched the families S24-7 (unclassified genus), Coriobacteriaceae (unclassified genus), Bacteroidaceae, as well as the genera *Bilophila* and *Bacteroides* relative to HF diet (**Figure S2D**). Importantly, we observed FFVHC diet promoted an early and significant expansion of *Bilophila* relative to both LF and HF diets (P<0.05) with a significant decrease in the genus *Bifidobacterium* relative to LF-fed animals (P<0.001) prior to the overt onset of disease at 8 weeks (**Figure S2E**). Despite no evident changes in the genera *Akkermansia* and *Dehalobacterium* after only 8 weeks of feeding, we noted that the genus *Sutterella* was also significantly increased by FFVHC diet compared to all other groups at this early time point (**Figure S2E**). These results indicate that increased dietary cholesterol in the FF diet results in a more rapid and potentially harmful gut dysbiosis that precedes disease onset, which could be possible indicators of disease progression and severity.

### Fast food diet containing high cholesterol induces gut dysbiosis that significantly correlates with clinical markers of NAFLD that predict outcomes

To determine whether gut microbiota imbalances induced by FF diets with increased cholesterol level exhibited associations with clinical markers in early vs. late disease, we performed Spearman correlation analysis between cecal gut microbiota features and alanine aminotransferase (ALT) levels in peripheral plasma of SPF mice at 8 and 24 weeks. Of the microbial features identified via LefSe analysis enriched by FF diets, we observed that both the genera *Bilophila* (P=0.000, r = 0.715), *Dehalobacterium* (P=0.041, r = 0.440), and *Bifidobacterium* (P=0.002, r = -0.662) were strongly and significantly correlated with plasma ALT levels at 24 weeks; both *Bilophila* and *Dehalobacterium* exhibited strong positive associations with plasma ALT while *Bifidobacterium* exhibited a strong negative association (**Figure 3A**). To determine whether these associations were also evident at early time points in disease course, we examined correlations between these ASVs and host ALT at 8 weeks. Importantly, we noted a strong and significant negative correlation between the relative abundance of the genus *Bifidobacterium* (P=0.015, r = -0.596) and host plasma ALT levels at 8 weeks (**Figure 3B**). However, neither *Bilophila* (P=0.484, r = 0.188) nor *Dehalobacterium* (P=0.996, r = -0.003) significantly correlated with host ALT levels at this early time point prior to disease onset. These data suggest loss of specific gut microbiota features, such as *Bifidobacterium*, induced by increasing levels of dietary cholesterol may be indicative of early disease, while an expansion of other community members, such as *Bilophila* may correlate with elevated inflammation during later disease stages.

**Figure 3.**
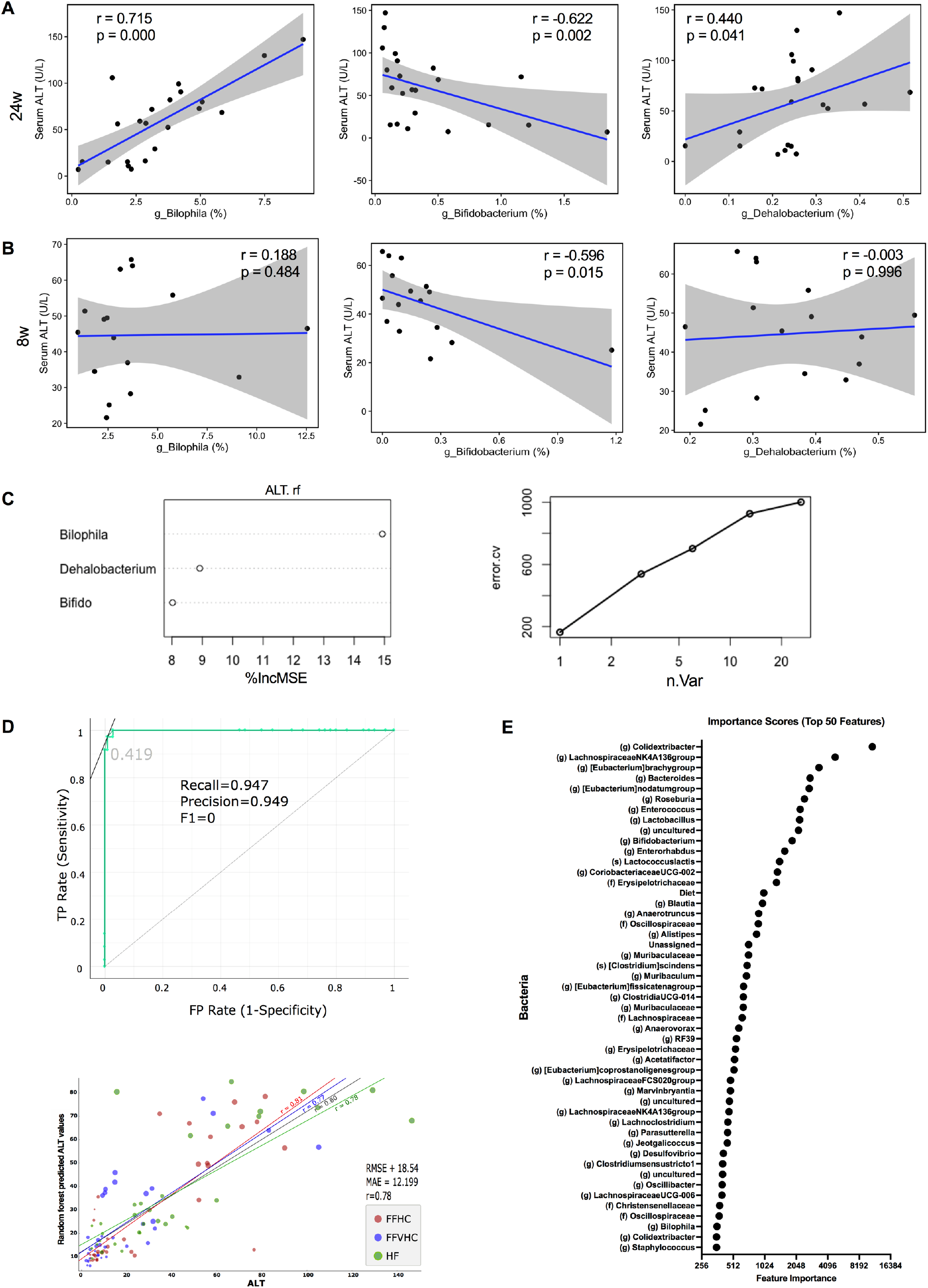
Fast food diet-induced gut dysbiosis significantly correlates with circulating levels of alanine aminotransferase (ALT) in peripheral plasma of SPF mice. (**A, B**) Spearman correlation analysis (p <0.01) between ALT and significantly shifted genera induced by fast-food diets identified by LEfSe at 24w (**A**) and 8w (**B**). r = correlation coefficient, p = p-value; 95% confidence intervals are shaded in gray. (**C**) Variable contribution and plot of general numbers vs. error rates. (**D**) The receiver operating characteristic (ROC) curve of the random forest (RF) model was constructed, and model specificity and sensitivity were tested by converting the predicted and actual ALT values into binned values of LOW, MEDIUM, and HIGH before evaluation (upper panel). AUC: 0.977, sensitivity or recall: 0.947, FP rate (1-sensitivity) or precision = 0.949. “TP: true positive rate”, “FP: false positive rate”, “Precision”, “Recall” and “F0”, “0.419” in the figure (lower panel) linear regression between predicted ALT scores from the RF model and actual ALT score for HF, FFHC and FFVHC groups. RMSE (root mean square error), mean absolute error (MAE), and the correlation between predicted and actual ALT score (r). (**E**) Feature scores generated from RF models to identify the top 49 taxa by plotting the increase in mean error rate when taxa were removed from the decision tree. The greater the Gini indices, the higher the importance of the variables. Diet was also selected as a feature of importance in the RF model.

Next, we examined whether late vs. early changes in gut microbiota community membership induced by FF diets detected via 16S rRNA gene amplicon sequencing in either cecal contents or feces could predict clinical disease outcomes, specifically host plasma ALT. Here, we constructed a model using Random Forest (RF) supervised machine learning. Initially, we generated a model for predicting ALT levels using the three significantly correlated genera, *Bilophila, Bifidobacterium*, and *Dehalobacterium* at 24 weeks (**Figure 3C**, left panel). The inclusion of these three genera explained 44.5% of the variation in ALT levels across all diets (**Figure 3C**, right panel). We next generated RF models to discern the importance of early changes in microbial taxa (4 and 8 weeks) in relation to plasma ALT levels later in disease (24 weeks) using ASVs in fecal samples collected over time (**Figure 3D**). The performance of the RF classification model based on fecal microbial ASV abundances sampled at 4 and 8 weeks resulted in an area under the ROC curve (AUC) of 0.977 in predicting ALT levels at 24 weeks for the validation set, corresponding to a sensitivity (recall) of 0.947 and FP Rate (1-Sensitivity, Precision) of 0.949 (**Figure 3D**, top panel). We noted a strong positive linear correlation between predicted ALT levels determined via the RF model and actual ALT levels for HF (r = 0.78), FFHC (r = 0.81) and FFVHC (r = 0.77) groups as well as across the entire dataset (r = 0.80) (**Figure 3D**, lower panel). The RF model selected 50 ASVs together with diet as the most important features that were predictive of ALT levels during disease onset and progression (**Figure 3E**). These 50 features were determined at the feature elimination step, and the best-performing model was selected as the final model. While several top features with the highest importance scores in the model using fecal ASVs over time were unique relative to those observed in cecal contents, we observed that the genera *Bifidobacterium* and *Bilophila* were in the top 50 (**Figure 3E**). Together, these findings indicate FF diet-induced gut microbiota imbalances, regardless of cholesterol level, are strongly associated with ALT levels in peripheral plasma of SPF mice. Further, early diet-induced fecal microbiota profiles are highly predictive of ALT levels later in disease, underscoring the potential of using feces coupled with dietary intake to serve as a non-invasive test to monitor NAFLD disease activity.

### Fast food (FF) dietary cholesterol levels and the presence of gut microbes enhance both local hepatic and systemic inflammation throughout the course of disease development

We next examined indicators of microbial translocation, specifically plasma lipopolysaccharide-binding protein (LBP), in SPF and GF animals. After 24 weeks, SPF mice fed FF diets, regardless of added cholesterol amount, exhibited significantly increased plasma LBP levels relative to LF-fed SPF counterparts, while, as expected, no differences were observed across groups in GF conditions (**Figure 4A**, left panels). While LBP was elevated in SPF mice at 8 weeks relative to GF, no differences across diets were detected regardless of microbial status (**Figure 4A**, right panels). These results suggest elevated LBP occurs later in disease progression after changes in gut microbiota are apparent and confirmed that increasing FF diet cholesterol level induces gut microbiota shifts that could induce systemic inflammation.

**Figure 4.**
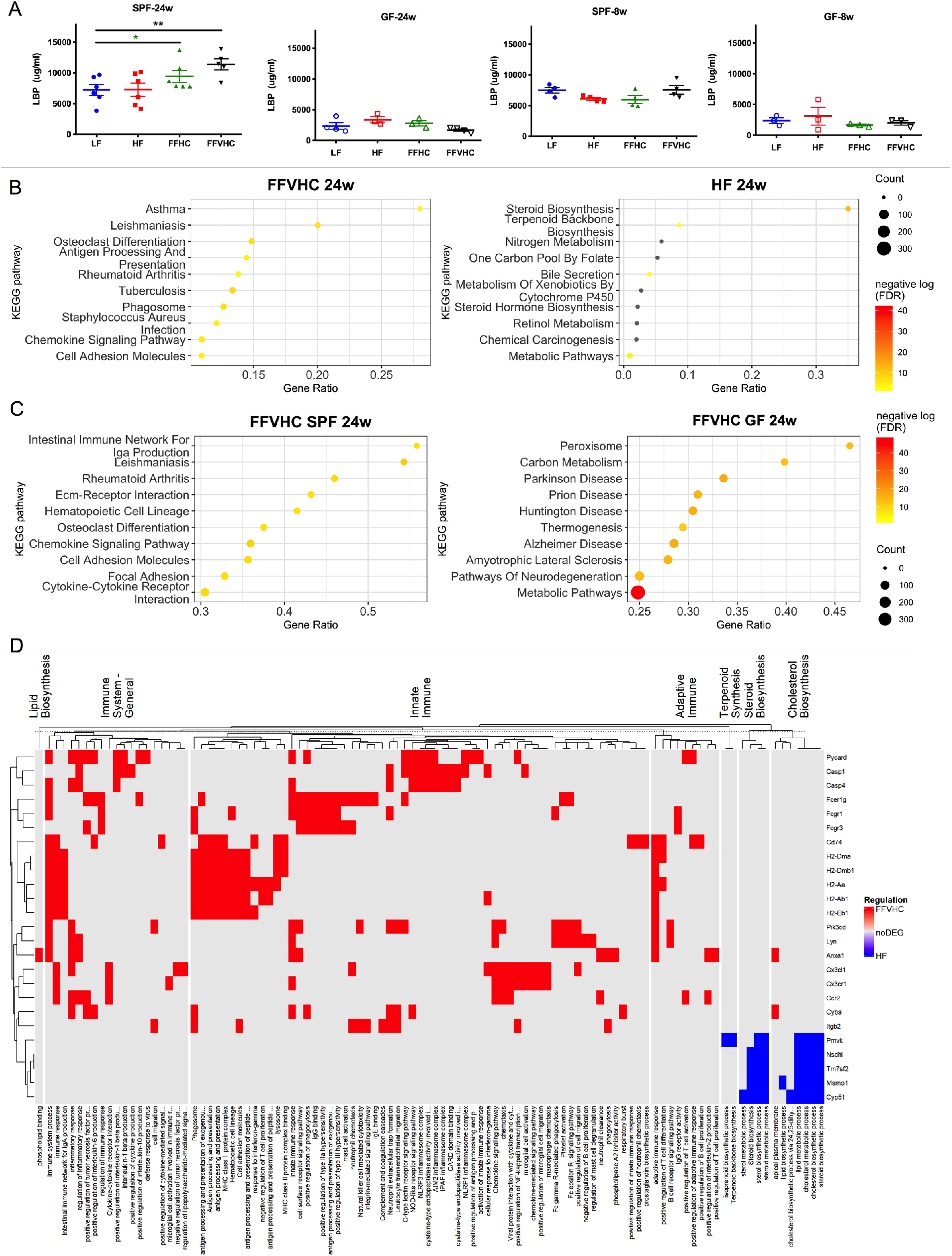
Dietary cholesterol level differentially impacts systemic and local inflammation in specific-pathogen-free (SPF) but not germ-free (GF) mice after 24 weeks of feeding. (**A**) Lipopolysaccharide binding protein (LBP) levels in peripheral plasma over time in SPF and GF mice. (**B, C**) Differentially expressed genes (DEG) sorted by the top 10 enriched categories from the background mouse genome in the hit ratio bar plot between SPF FFVHC-vs. SPF HF-fed mice (**B**) and between SPF FFVHC-vs. GF FFVHC-fed mice (**C**) after 24 weeks of feeding. (**D**) Heatmap of enriched GO terms and KEGG pathways grouped into major categories based on immune and metabolism-related functions. Up- and down-regulated DEGs in the grouped GO terms or KEGG pathways are distinguished by red and blue, respectively. n=4-6 per group. Data represent means± S.E.M. *P <0.05 via nonparametric Kruskal-Wallis test, followed by Turkey’s multiple comparisons between the means of all groups. Star color represents the treatment group exhibiting significant differences.

To explore this, we next performed RNA sequencing to characterize how diet, as well as the presence or absence of gut microbiota, influence local liver inflammation after both 8 and 24 weeks. Significant gene expression differences between GF and SPF mice across diets were identified using log2 fold change and FDR adjusted p-value (p<0.05). Given our observations that FFVHC appeared to elicit the greatest impact on both host and microbe phenotypes, we focused our analyses on comparisons between HF and FFVHC under GF and SPF conditions to home in on the impact of very high cholesterol exposure. We performed enrichment analysis of significant differentially expressed genes (DEGs) with mouse-specific GO terms and KEGG pathways. Here, after 24 weeks, the top 10 enriched KEGG pathways (category 3) in livers from SPF FFVHC-fed mice exhibited significant and differential enrichment for immune-related pathways relative to HF-fed counterparts, including those associated with asthma, leishmaniasis, osteoclast differentiation, antigen processing and presentation, Rheumatoid Arthritis, tuberculosis, phagosome, staphylococcus aureus infection, chemokine signaling pathways, and cell adhesion molecules (**Figure 4B**, right panel). Conversely, KEGG pathways containing DEGs involved in steroid biosynthesis, terpenoid backbone synthesis, bile secretion, and global metabolic pathways were significantly downregulated in FFVHC-fed SPF mice (**Figure 4B**, right panel). Interestingly, after only 8 weeks of FFVHC feeding, significant downregulation of DEGs in nearly identical KEGG pathways was observed, including steroid biosynthesis, fatty acid elongation, biosynthesis of unsaturated fatty acids, and terpenoid backbone biosynthesis, fatty acid metabolism, peroxisome, and metabolic pathways relative to HF-fed mice (**Figure S4A**). However, at 8 weeks, no significant pathway enrichment was observed in FFVHC relative to HF-fed conditions. Together, these data suggest immune dysregulation induced by high saturated fat coupled with elevated cholesterol intake occurs following diet-mediated shifts in gut microbiota and much later in disease progression.

We next examined enriched KEGG pathways determined via DEGs in the liver between FFVHC-fed SPF vs. GF mice to identify contributions of gut microbiota in the context of cholesterol-rich diets and NAFLD development at 8 and 24 weeks. Here, we observed the majority of the top 10 significantly enriched KEGG pathways from DEGs up-regulated in FFVHC-fed SPF vs. GF mice after 24 weeks were those associated with immune function (**Figure 4C**, left panel), confirming gut microbes are required for upregulation of inflammation. These pathways included IgA production, leishmaniasis, Rheumatoid Arthritis, Extracellular matrix receptor interaction, hematopoietic cell lineage, osteoclast differentiation, chemokine signaling pathways, and cell adhesion molecules. Conversely, FFVHC-fed SPF mice exhibited downregulation of pathways related to metabolism, including Peroxisome, Carbon metabolism, Parkinson’s disease, Prion disease, and thermogenesis relative to GF FFVHC-fed mice (**Figure 4C**, right panel). At 8 weeks, we noted livers from FFVHC-fed SPF animals exhibited enrichment in pathways associated with oxidation, including Proteasome, Glutathione metabolism, and metabolism of xenobiotics via cytochrome P450, and several immune pathways, particularly phagosome and Salmonella infection (**Figure S4B**). However, GF FFVHC-fed mice exhibited pathway enrichment in amino acid metabolism, including arginine biosynthesis, glycosaminoglycan biosynthesis, and lysine degradation. Finally, examining changes over time only in FFVHC-fed SPF mice, we noted several immune-related pathways were significantly enriched based on elevated DEGs after 24 weeks relative to 8 weeks (**Figure S4B**). These data underscore gut microbes are a prerequisite for FFVHC diet-induced NAFLD onset and progression, which contributes to inflammation in advanced disease.

To further assess the role of HF vs. FFVHC in driving immune versus metabolic functions in SPF animals, we grouped enriched GO terms and KEGG pathways into these two major categories. As shown in **Figure 4D** (heatmap), livers from 24-week SPF FFVHC-fed mice exhibited upregulation of immune-related genes across general, innate, and adaptive immune pathways responsible for phospholipid signaling, pro-inflammatory cytokine production, inflammasome activation, antigen-processing and presentation, as well as cytokine and chemokine signaling relative to HF. Conversely, HF diet led to upregulation of genes involved in lipid metabolism pathways, particularly biosynthesis of terpenoids, steroids, and cholesterol relative to FFVHC. After only 8 weeks of diet, there was no indication of upregulation of immune-related programming in the livers of FFVHC-fed SPF mice (**Figure S4C**). Surprisingly, after 8 weeks, FFVHC diet upregulated genes involved in biosynthesis of lipids, cholesterol, steroids, and terpenoids relative to HF. These data underscore the significant contribution of gut microbes to FF diet-induced upregulation of immune-associated pathways in the liver, specifically later in the disease. Taken together, these results indicate gut microbes are required for diet-induced inflammation in NAFLD to NASH transition, i.e., GF mice are protected against disease, and that levels of dietary cholesterol in conjunction with exposure to high fat and high sugar can profoundly accelerate inflammation in the presence of a complex microbial community.

### Increased cholesterol level in fast food diets exacerbates gut microbiota functional imbalances that precede onset of NAFLD

Given our observations that shifts in gut microbiota community membership occur rapidly in response to diet, are dependent on dietary cholesterol level, serve as predictive markers of disease, and contribute to the host’s pro-inflammatory status, we next examined fecal gut microbiome functional properties via shotgun metagenomics in 4 mice per treatment group after 8 and 24 weeks of diet exposure. Since we used semi-purified diets that contain identical basal ingredients, we did not anticipate overt global changes to microbial metabolic profiles; instead, this design would allow us to observe shifts in specific functions. After 24 weeks, as anticipated, PCoA of short sequencing reads revealed no significant differences in general metabolic pathway abundances between all groups (**Figure S5A**), which is also evident in the heatmap (**Figure S5B**). However, LEfSe analysis indicated the abundances of several pathways differed in FFHC and FFVHC-fed mice relative to HF-fed controls (LDA score (log10) ± 3 or more; **Figure S5C**). For instance, FFVHC differentially upregulated methane metabolism, cell cycle (ko04112), Pentose phosphate pathway (ko 00030), RNA polymerase (ko 03020), as well as taurine and hypotaurine metabolism (ko 00430) (**Figure S5C**). Conversely, HF diet enriched pathways associated with C5 Branched dibasic acid metabolism (ko 00660), Terpenoid backbone biosynthesis (ko 00900), Histidine metabolism (ko 00340), and RNA degradation (ko 03018) relative to FFVHC (**Figure S5C**). Interestingly, HF feeding appeared to upregulate LPS biosynthesis relative to FFHC; however, circulating plasma LBP remained low relative to FF diets, indicating additional factors may contribute to gut barrier dysfunction upon incorporation of increasing dietary cholesterol.

To characterize metabolic feature distribution across taxonomic profiles identified via MetaPhlAn, we applied HUMAnN to 8- and 24-week fecal samples. The resulting organism-specific gene hits were functionally assigned to pathways using MinPath, and their relative abundances were assessed. We observed significant differences based on stratified contributions of functions attributed to key taxa within fecal samples from FFVHC-fed mice. This revealed FFVHC diet reduced levels of potentially beneficial functions attributed to microbes associated with reduced gut inflammation relative to both LF and HF-fed controls after 8 and 24 weeks (**Figure 5** and **Figure S5D-H**). For example, after 24 weeks, we observed reductions in functions associated with carbohydrate metabolism (**Figure 5A**), amino acid metabolism (**Figure 5B**), and metabolism of vitamins (**Figure 5E**) attributed to *Bifidobacterium psuedolongum* and *Akkermansia muciniphila* species. Conversely, FFVHC feeding increased pathways associated with specific species that may contribute to gut inflammation. For instance, *Parasutterella excrementihominis*, which was elevated in FFVHC, exhibited increases in pathways associated with carbohydrate metabolism (**Figure 5A**), amino acid metabolism (**Figure 5B**), lipid metabolism (**Figure 5C**), glycan biosynthesis and metabolism (**Figure 5D**), and metabolism of vitamins (**Figure 5E**). A similar trend was observed as early as 8 weeks after diet exposure (**Figure S5D-H**).

**Figure 5.**
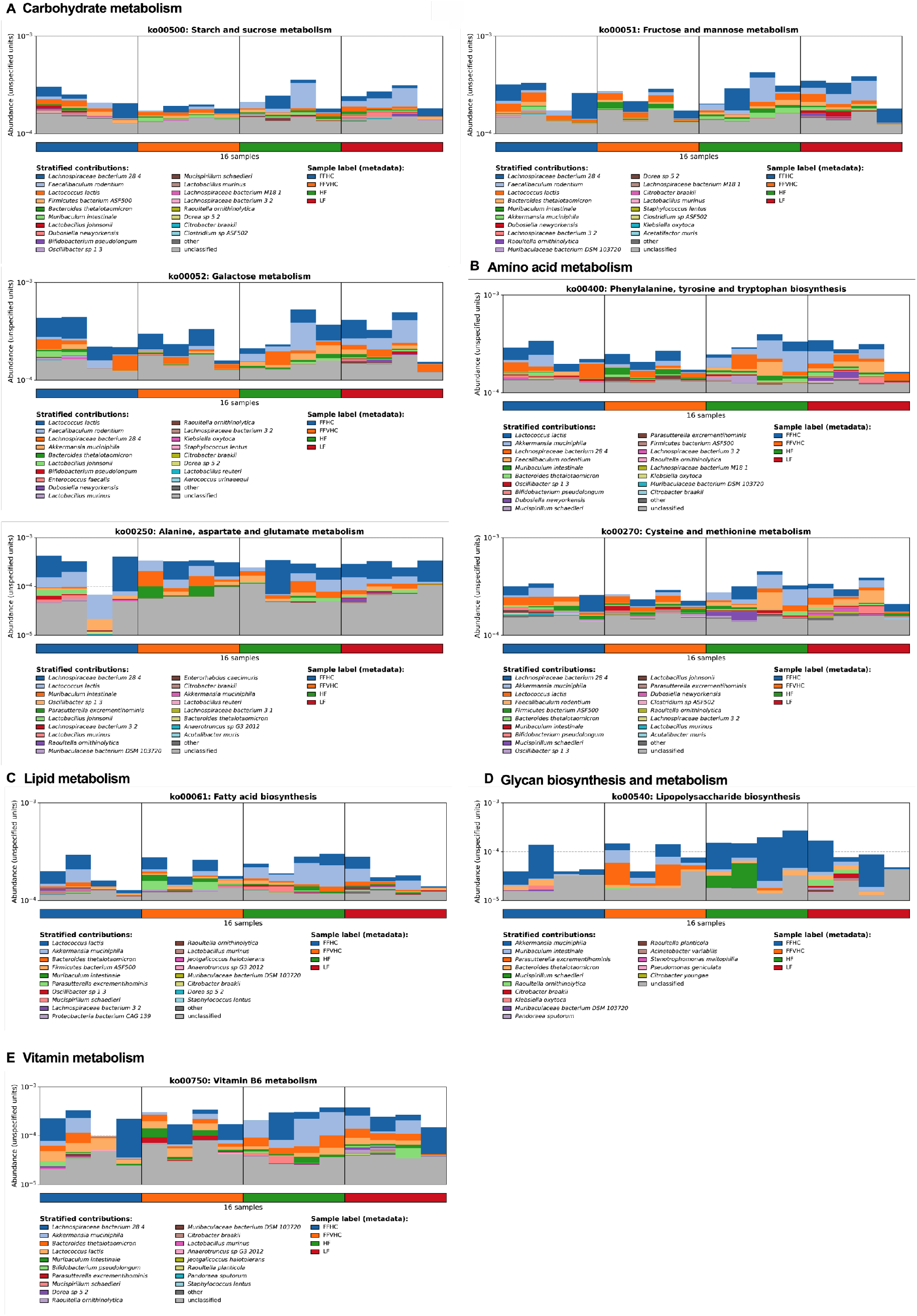
Fast food diet promotes gut microbiota functional dysbiosis in feces that corresponds with NAFLD outcomes after 24 weeks of feeding. (**A-E**) Microbiome-derived metabolic features differentially induced by diet separated based on MetaPhlAn taxonomic profiles indicating shifts in (**A**) Carbohydrate metabolism, (**B**) Amino acid metabolism, (**C**) Lipid metabolism, (**D**) Glycan biosynthesis and metabolism, and (**E**) Metabolism of vitamins.

### Fast food diets containing high cholesterol reduce network co-occurrence patterns amongst gut microbes and increase overall community vulnerability to subsequent perturbations

To examine how FF diets containing increasing cholesterol influence overall gut microbiota community networks, co-occurrence analysis was performed via Spearman correlation between 16S rRNA gene ASVs in cecal contents collected from mice after 24 weeks on diets. Only robust correlations were included for inferred network construction (inclusion criteria: -0.60 ≥ ρ ≥ 0.60, FDR p <0.05) (**Figure 6**). Regardless of diet, the inferred networks were dominated by the two major intestinal phyla, Firmicutes and Bacteroidetes, with decreased contributions from less abundant community members. We noted that diet composition significantly impacted network connectivity; the inferred network for LF diet exhibited a high number of nodes (218) and edges (918) compared to all other groups. Surprisingly, the addition of fat alone (HF) resulted in a significant increase in edges (258 nodes and 5349 edges; **Figure 6A**). However, upon addition of dietary cholesterol, we observed a significant decrease in both nodes and edges in FF diets regardless of added cholesterol level (FFHC = 174 nodes, 742 edges: FFVHC = 160 nodes, 644 edges) relative to both LF and HF (**Figure 6A**).

**Figure 6.**
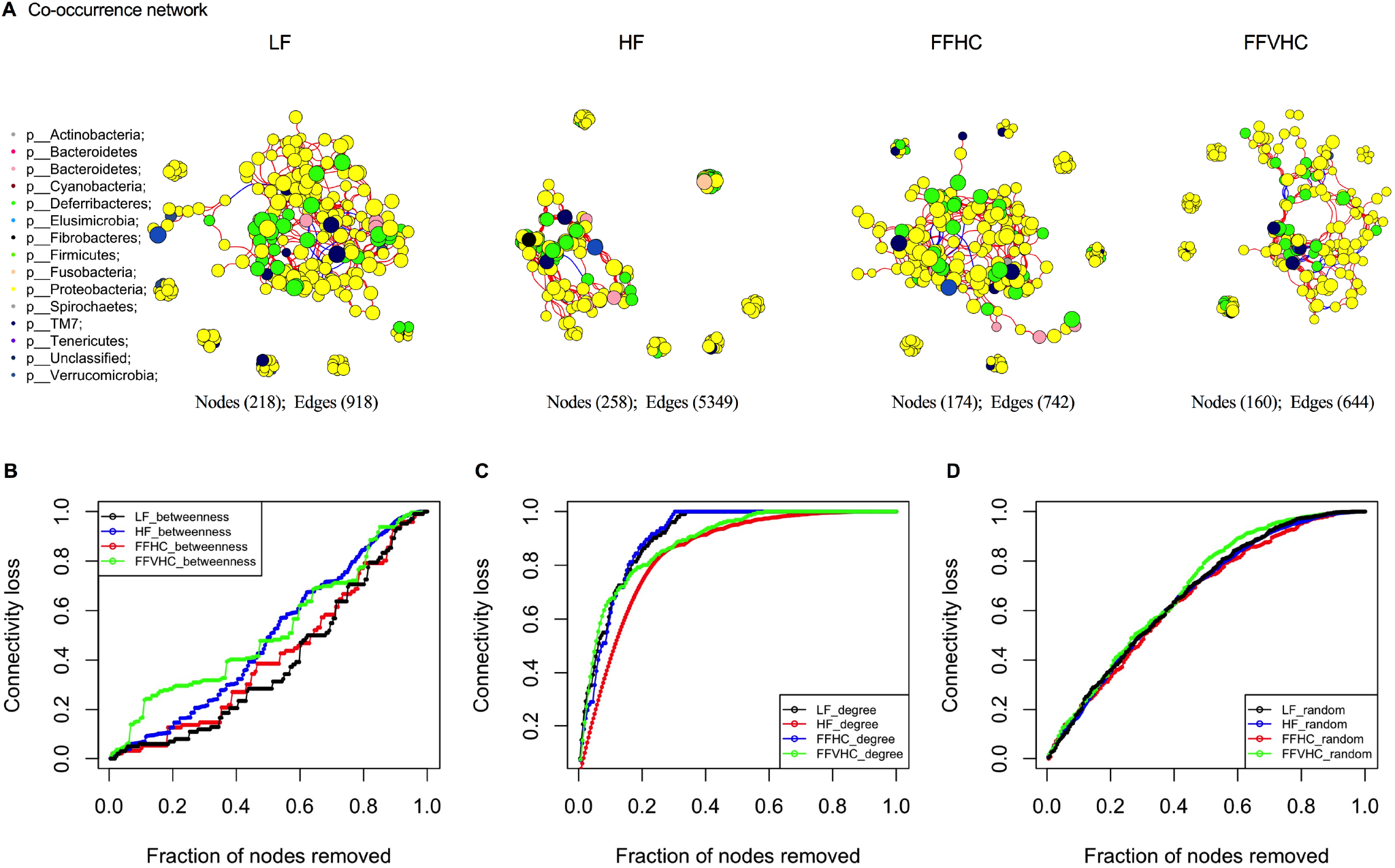
Network analysis reveals less resistant and unstable co-occurrence patterns in gut microbes among mice fed FF diets containing high cholesterol after 24 weeks. A connection represents a strong (Spearman correlation coefficient of *ρ*>0.6 and significant (*P*<0.05) correlation. **A**, gut microbiota co-occurrence networks from mice fed LF diet, HF diet, FFHC diet, and FFVHC diet after 24 weeks. The vertices represent unique ASV features in each data set. The size of each vertex is proportional to the relative abundance of each ASV. Vertex color corresponds to phylum taxonomic classification. The edge thickness is equivalent to correlation values. (**B-D**) Error and attack tolerance tests were performed stepwise by removing vertices from the network in decreasing order of betweenness (**B**), closeness (**C**) and randomly (**D**) and recording the network’s loss of connections (edges).

We next tested resilience of the inferred networks induced by diet via systematic removal of ASVs in decreasing order of betweenness (a measure of the number of shortest paths going through a vertex), closeness (the steps required to access every other vertex from a given vertex) or randomly, which indicates the microbial community’s resistance to or potential loss of connectedness upon perturbation. To assess network robustness, we performed ASV extinction simulations using betweenness scores. Here, key ASVs were removed stepwise from the network in decreasing order of betweenness and loss of connectivity was recorded. Targeted removal of vertices with the highest betweenness centrality from the co-occurrence network of LF-fed mice led to a gradual loss of connectivity (∼20%) after removing 40% of the vertices (**Figure 6B**), indicating the LF diet-induced microbial community exhibits a high buffering capacity. This network only becomes compromised after removal of a high percentage of the most central community members, where a total breakdown (loss of 95% of the connectivity) was observed after removing the top 90% of vertices with the highest betweenness values. Conversely, gut microbiota co-occurrence networks from HF, FFHC, and FFVHC-fed mice were more vulnerable and less robust upon ASV removal (**Figure 6B**), particularly FFVHC diet relative to LF controls. The rapid and persistent loss of connectivity in FFVHC co-occurrence networks upon initial and subsequent ASV removal suggests greater microbial community vulnerability (up to 28%, 12.5%, and 29% loss after removing 40% vertices for HF, FFHC and FFVHC, respectively, **Figure 6B**). The total breakdown of community networks in mice fed HF or FFVHC diet was achieved earlier than in mice fed LF diet (top 85% vertices for HF, top 89% vertices for FFHC, and top 84% vertices for FFVHC, respectively). Surprisingly, total breakdown of the co-occurrence network of FFHC was achieved at removal of the top 89% of vertices, which was approximately equal to that of LF (**Figure 6B**). These data indicate FF diets containing elevated cholesterol lead to gut microbiota network instability which could be linked, in part, to a loss of functional elements from key gut microbiota community members. Targeted removal of vertices with the degree or randomly both showed rapid decrease of network robustness and breakdown of network structure (**Figure 6C and D**). These alterations in connectivity and susceptibility to perturbation were not simply due to reduced alpha-diversity in FF diet-fed animals, as no significant differences in this metric were observed (p>0.05, data not shown). These data indicate FF diets containing elevated cholesterol drive gut microbiota network instability which could be linked, in part, to a loss of functional elements from key gut microbiota community members.

## Discussion

While previous studies suggest gut dysbiosis is associated with NAFLD/NASH development in patient populations, the underlying etiology of human disease is difficult to recapitulate in mice. Here, using the murine FF diet model, we demonstrate gut dysbiosis is evident in stool after only 4 weeks of FFVHC diet, preceding overt hepatic disease onset and serving as a highly predictive marker of histological progression of NAFLD/NASH. In mice harboring gut microbes, dietary cholesterol level exacerbates both gut dysbiosis and hepatic inflammation, while GF mice are protected from FF diet-induced NAFLD/NASH, identifying gut microbes as a prerequisite for disease. Using this model system, our studies provide the first insight into the timing and interactions between complex Western-style diets containing high cholesterol and gut microbes in NAFLD/NASH onset, development, and progression.

Previous studies have qualitatively shown the microbiome rapidly shifts due to diet, but long-term consequences to host health are poorly understood (Sonnenburg and Bäckhed 2016). To drill down on specific dietary components in the context of NAFLD and gut dysbiosis, we utilized semi-purified diets for both LF control as well as in our HF and FF diets, expanding on previous studies employing this model, where undefined chow diet has been utilized as the control (Charlton et al. 2011, Krishnan et al. 2017). Here, LF, HF, FFHC, and FFVHC diets contained identical ingredients, including fat source (anhydrous milkfat, **Table 1**). Initial shifts in gut microbes from baseline to 4 weeks were mainly explained by the shift from standard chow to semi-purified diets, which showed large alterations in bacterial composition with the same trend across all groups (**Figure S2A**), identical to what others have shown using similar diets (Devkota et al. 2012, Huang et al. 2013). We show early changes in microbiota community membership and function occur before NALFD histological onset and that by week 24, FFVHC-fed mice exhibited increased liver steatosis with more severe NAS score as compared to both LF and HF-fed mice, recapitulating previous findings (Charlton et al. 2011, Krishnan et al. 2017). Molecular indicators of genes involved in fibrosis identified via RNA sequencing of liver, including α-smooth muscle actin (*ASMA*), type I collagen (*Col1a1*), transforming growth factor beta 1 gene (*Tgfb1*), hepatocyte growth factor (*Hgf*), Tissue inhibitor of metalloproteinase-1 (*Timp1*), and *Lumican* in SPF FFVHC-fed mice relative to LF-fed counterparts indicated trends toward a transition to fibrosis (data not shown). The early microbial shifts include a loss of key members, particularly *Bifidobacteria*, after only 4 weeks on FFVHC diet (data now shown), which is also evident in both feces and cecum after 8 weeks of feeding prior to disease onset. Depletion of *Bifidobacteria* is further evident by increasing dietary cholesterol intake (**Figure 2D**), and decreased relative abundance of this genus negatively correlates with exacerbated biomarkers of liver injury, e.g., ALT, which remains the best clinical predictor of disease severity and outcome (Thapa and Walia 2007)) throughout the course of disease. Later in disease progression, pro-inflammatory microbiota indicators emerge, i.e., *Bilophila*, which could be critical inducers and drivers of host inflammation, contributing to NAFLD to NASH transition. This expansion of *Bilophila* late in disease also positively correlates with host ALT, but this is not observed early in disease course (**Figure 3A** and **B**). Our results are consistent with those of previous studies regarding the relationship of these bacteria with liver diseases. For example, the abundance of *Bifidobacterium* has been reported to decrease in individuals with NAFLD/NASH (Leclercq et al. 2014, Xu et al. 2012), while another study showed that probiotic *Bifidobacterium spp*. delivery could attenuate hepatic injury (Liu et al. 2019). Further, the genus *Bilophila* has been found to be enriched in diet-induced NAFLD/NASH, which promotes a pro-inflammatory response (Natividad et al. 2018). These critical members of the gut microbiota community can be impacted by HF, high cholesterol, and high simple sugar diets (Natividad et al. 2018, Zhang et al. 2021), which can contribute to complex liver disease. Recent studies also identified numerous gut microbiota that can metabolize cholesterol and steroidal compounds, providing insights into the molecular mechanisms by which microbial community members modify broad classes of dietary lipids, which alters host uptake (Le et al. 2022, Yao et al. 2022). Interestingly, Le et al., showed that *Bifidobacterium psuedolongum* exhibited the highest capacity for exogenous cholesterol uptake in mice harboring a complex microbial community. Whether and how extreme levels of cholesterol modify this interaction remains to be determined and further investigations are needed. Together, our results indicate FF diets containing high cholesterol result in a rapid induction of “harmful” gut dysbiosis with loss of key community members that may contribute to earlier NAFLD onset and progression.

**Table 1.** Diet Composition used in the animal trial.

| Diets | Chow | LF* | HF* | FFHC* | FFVHC* |
| --- | --- | --- | --- | --- | --- |
| Protein % | 19.8 | 19.1 | 15.1 | 15.2 | 15.4 |
| Carbs % | 54.1 | 67.9 | 43.1 | 43 | 42.2 |
| Fat % | 26.1 | 13 | 41.8 | 41.9 | 42.5 |
| Total Cholesterol % | 0.07 | 0.05 | 0.05 | 0.2 | 2 |
| Kcal/g | 3.6 | 3.6 | 4.6 | 4.6 | 4.5 |

\*LF = Low fat without added cholesterol, HF = High fat without added cholesterol, FFHC = High fat with 0.2% added cholesterol; FFVHC = High fat with 2% added cholesterol. %, percentage of weight.

Our study shows early gut microbiome changes induced by diet not only correlate with but can predict disease severity and progression. Our RF models demonstrate early diet-induced gut microbiota profiles are predictive of ALT levels late in disease, underscoring the potential of diet-induced gut microbiota as a non-invasive test to monitor NAFLD disease activity. Thus far, several studies have shown gut microbiota community membership and functional outputs can predict and categorize liver disease and fibrosis stage in NAFLD/NASH patients (Loomba et al. 2017, Caussy et al. 2019). Further, Kang et al. demonstrated that high, but not low, carbohydrate dietary intake was associated with an altered gut microbiome and clinical markers of NAFLD, also revealing the capacity of three high carbohydrate-induced bacterial families to predict NASH outcomes in a Korean population (Kang et al. 2022). Yet, due to the observational nature of this study not all dietary components could be controlled for, and the authors conceded that both low- and high-carbohydrate intake groups consumed relatively high amounts of carbohydrates. Xiang et al. identified that cholesterol-induced gut microbiota dysbiosis contributed to NAFLD– Hepatocellular carcinoma (HCC) development in mice (Zhang et al. 2021), however, in this study, the predictive role of gut dysbiosis corresponding to specific diet components remained unexplored. Here, we advance these findings in our model system, identifying that dietary composition is a key factor in driving gut dysbiosis (i.e., dietary cholesterol level) and, when considered in the model, can serve as a sensitive and key clinical indicator of disease. This requires validation in human populations that include deep phenotyping with dietary intake and gut microbiota read-outs as key elements. For instance, the NHANES study (Christensen et al. 2013) showed that the risk of liver disease-related mortality was dramatically increased in subjects taking in cholesterol greater than 511 g/day, however, no information on gut microbes is available for these subjects. Our work indicates dietary fat and cholesterol coupled with gut microbes are key drivers and should be considered in NAFLD/NASH disease development and severity.

Co-occurrence networks showed significant differences in the vertical-level features (i.e., vertices and edges) induced by different diets. Specifically, the FFVHC microbial community showed markedly fewer vertices and edges, indicating lower bacterial diversity and a simpler network relationship among community members. Targeted, intentional attacks via decreasing order of betweenness (Albert, Jeong and Barabási 2000) in FFVHC revealed decreased gut microbiota network robustness and resilience relative to all other diets, particularly LF. No significant differences in alpha diversity were detected across diets, suggesting that the tolerance difference was not related to reductions in diversity. It remains unclear from our studies whether early onset of gut dysbiosis contributes to gut microbiota community membership instability and the subsequent influence on disease outcomes, or if host-associated disease processes (i.e., upregulation of cytokines, shifts in bile acids, etc.) feed back onto microbial networks to induce disruption. Nevertheless, we reveal FF diet containing high cholesterol drives less robust co-occurrence patterns amongst gut microbes and increases the overall gut microbiota community vulnerability to perturbations, indicating reduced capacity to maintain host physiology and metabolic homeostasis.

In our study, FF dietary cholesterol level differentially impacts short- and long-term systemic and local inflammation in SPF, but not in GF mice. Collective evidence highlights NAFLD/NASH is characterized by chronic low-grade inflammation (‘metaflammation’) in animal and human studies (Furman et al. 2019, Hummasti and Hotamisligil 2010, Targher, Byrne and Tilg 2020), which is driven by gut microbiota (Carpino et al. 2020, He et al. 2021, Tilg and Kaser 2011). The endotoxin-producing gut bacteria from NAFLD/NASH individuals played a causal role in NAFLD development in GF mice fed high-fat diet (Fei et al. 2020). Here, we further confirmed an essential role for gut microbes in FF-diet-induced systemic inflammation regardless of cholesterol level. The local transcriptional pro-inflammatory signature in the liver determined via RNA sequencing, as well as that detected in the gut microbiota over time using metagenomic sequencing, also underscores significant contributions of microbiota, particularly those induced by FF diet containing high cholesterol on the transition from NAFLD to NASH. How dietary cholesterol interacts with high fat and high sugar to accelerate inflammation in the presence of a complex microbial community requires further dissection and should be a focus of future studies. Collectively, these findings aid in defining the timing and mechanistic insights into the pathways contributing to NAFLD/NASH development, dietary factors, gut dysbiosis, and immune dysregulation.

In conclusion, our principal findings reveal early gut microbiota dysbiosis (membership and function) induced by FF diets containing elevated cholesterol is highly predictive of NAFLD/NASH outcomes, suggesting gut dysbiosis is an initiator of metabolic liver disease. These findings may have profound clinical implications for the timing of dysbiosis in relation to transition to advanced disease stages, underscoring the pivotal role of dietary intake (specifically fat and cholesterol) as a central factor in these processes. First, by providing a time-course analysis of the gut microbiota, it is possible to identify microbially-derived biomarkers that predict and stratify individuals at high risk for development and aggressive progression of NAFLD/NASH. Knowing this – there is an opportunity to identify individuals who might best benefit from microbiome/diet-based interventions. Of all the organ systems in the body, the microbial organ is the most manipulatable, and metagenomic or metabolomic profiles can identify specific functional subsystems to correct to lower risk. Finally, these studies provide opportunities to identify specific taxa and metabolic mediators that can be developed as future therapeutics. Taken together, this work has set the framework for development of microbiome-based interventions that could potentially prevent or reverse NAFLD/NASH disease course, reducing overall mortality.

### Limitations of the study

While our findings provide insights into the time course and predictive element of diet-induced gut microbes, some limitations exist. First, we identified early onset of gut dysbiosis, less robust and unstable co-occurrence patterns, and immune dysregulation, however, the causal role for these early changes on later stage disease processes remains elusive. Whether early losses of or emergence of typically low-abundance community members later in disease and which taxa or functions are most important remain to be elucidated. Gnotobiotic animal studies with FMT using early vs. late dysbiotic communities induced by FF diets will be required to parse this out in future experiments. We recognize that cholesterol levels contained within the FFVHC diet is equivalent to eating 50-60 eggs per day, which is considered supraphysiological. While this level of cholesterol is less relevant to the general human population, it allowed for a deeper understanding of cholesterol’s interactive role in NAFLD onset and progression in the presence or absence of gut microbes. Despite these limitations, our study reveals an association between earlier onset of gut dysbiosis in FF-fed mice contributing to more severe disease. This study has set a framework from which to build on to provide further insights into the causation of dysbiosis induced by FF diets both in preclinical and clinical settings.

## ACKNOWLEDGMENTS

The authors are grateful to the University of Chicago Animal Resources Center and the Gnotobiotic Research Animal Facility for SPF and GF mouse husbandry as well as Nicholas Rinella for technical assistance. V.A.L. received support from the Gilead Research Scholars Program Foundation for Liver Diseases and NIH NIDDK K01DK111785. J.B.H. received support from NIH MANTP T32 DK007665. V.A.L., M.R.C., O.D. and N.F. received support from the Wisconsin Alumni Research Foundation, Accelerator Microbiome Challenge Grant. V.A.L., M.R.C., and E.B.C. received support from the University of Chicago Gastrointestinal Research Foundation.

## AUTHOR CONTRIBUTIONS

V.A.L., M.R.C., and E.B.C directed and designed the study. N.F., S.M., J.M., O.D., M.H., J.B.H., W.C., J.H., and V.A.L. managed and performed the mouse work, sample analyses, and data interpretation. N.F. and O.D. analyzed and interpreted the microbiota data; B.X., M.D., and D.S. analyzed RNA sequencing data; N.F., K.B.M-G., V.A.L., and M.R.C. wrote the paper. All authors reviewed and approved the final manuscript.

## Materials and Methods

### Animals, Diets, and Experimental Timeline

Specific pathogen-free (SPF) litter-matched eight-week-old male C57Bl/6J mice were purchased from Jackson Laboratories. All the mice were maintained on a 12:12-h lighted-dark cycle. Upon arrival, mice were group-housed on pine shavings (*n*=4/cage) and acclimatized within the University of Chicago SPF barrier facility for 8 weeks. Here, they received bi-weekly mixed bedding as previously described to minimize cage effect and provided *ad libitum* access to standard Purina LabDiet® 5K67 autoclaved rodent chow (Table 1).

Germ-free (GF) male C57BL/6 mice were bred under GF conditions at the University of Chicago Gnotobiotic Research Animal Facility, group-housed on pine shavings and provided *ad libitum* access to standard Purina LabDiet® 5K67 autoclaved rodent chow (Table 1). The GF status of the mice was assessed every week using aerobic and anaerobic cultivation and 16S rRNA PCR from feces to confirm the absence of bacteria, molds, and yeast throughout the experiment.

At 16 weeks of age, SPF or GF mice were housed 2/cage (week 0) for 8 or 24 weeks, as previously reported (Charlton et al. 2011, Krishnan et al. 2017). While previous comparisons using the FF model relied on grain-based, undefined chow-fed controls, this study utilized a semi-purified low-fat diet to avoid potential confounders of diet ingredients. All purified diets were customized and purchased from Envigo. Mice were randomly assigned to one of four diets: (1) low-fat diet without added cholesterol (LF), (2) high-fat diet without added cholesterol (HF), (3) fast food (same fat content as HF diet group) with 0.2% added cholesterol (FFHC) or (4), modified fast-food with 2% added cholesterol (FFVHC) (Table 1). All mice received drinking water containing fructose (23.1 g/l) and glucose (18.9 g/l). Mice were tracked and sacrificed after 8 or 24 weeks of dietary exposure, consistent with previous studies (Charlton et al. 2011, Krishnan et al. 2017). During the tracking period, weekly body weight, water consumption, and diet consumption were assessed, and monthly fecal and blood samples were collected. Mice were euthanized using CO_2_, and tissue samples were harvested at the end of the trial. At 8- or 24-weeks, livers were flash-frozen and stored at -80°C for subsequent assays. Adipose tissues (mesenteric, gonadal, inguinal, and retroperitoneal) were weighed, and the remaining tissues were flash-frozen for future analysis.

### Serum biochemical analysis

To examine liver function, a colorimetric-based enzyme assay was used to measure alanine aminotransferase (ALT) levels in peripheral plasma collected from the submandibular vein monthly using a Mouse ALT Activity Assay Kit (SIGMA). Serum LBP level was measured by a Mouse Lipopolysaccharide Binding Protein ELISA Kit (HyCult Biotechnology, Uden, The Netherlands). All assays were performed according to the manufacturer’s instructions.

### Liver histology

Liver samples were cryopreserved in OCT media, stored at -80°C, and sectioned using a cryotome. Frozen liver sections were stained with Oil-Red-O. Additionally, a separate liver sample from each mouse was formalin-fixed, embedded in paraffin, and sectioned (5 μm) followed by deparaffination for hematoxylin-eosin (H&E) and Masson’s trichrome staining using standardized protocols, and scored for severity of steatosis and inflammation by an experienced pathologist blinded to treatment. A semi-quantitative system of scoring NAFLD features (NAFLD Activity Score [NAS]), was developed based on histological assessment of H&E staining. Total NAS, ranging from 0-8, represents the sum of scores for steatosis (0–3), lobular inflammation (0–3), and ballooning (0–2). Steatosis scores: percentage of surface area containing steatosis as evaluated on low to medium power examination; 0 (0–5%), 1 (5–33%), 2 (33–66%), 3 (>66%). Lobular Inflammation scores: 0 (None), 1 (< 2 foci / 200x), 2 (2-4 foci / 200x), 3 (> 4 foci / 200x). Hepatocyte Ballooning scores: 0 (None), 1 (Few balloon cells), 2 (Many cells/prominent ballooning) (Kleiner et al. 2005).

### DNA isolation and 16S ribosomal RNA (rRNA) gene sequencing

DNA was extracted from fecal samples collected monthly using standard, published protocols (Miyoshi et al. 2018). The V4 region of the 16S rRNA gene was amplified, and sequences were obtained by Illumina MiSeq via the Next Generation Sequencing Core in the Biosciences Division at Argonne National Laboratory (Walters et al. 2016). The primers used for amplification (515F-806R) contain adapters for MiSeq sequencing and single-end barcodes allowing pooling and direct sequencing of PCR products (Caporaso et al. 2012). Each 25 μl PCR reaction contained the following mixture: 12 μl of MoBio PCR Water (Certified DNA-Free; Mo Bio Laboratories), 10 μl of 5-Prime HotMasterMix (1×), 1 μl of forward primer (5 μM concentration, 200 pM final), 1 μl of Golay Barcode Tagged Reverse Primer (5 μM concentration, 200 pM final), and 1 μl of template DNA (Leung et al. 2018). The conditions for PCR were as follows: 94 °C for 3 min to denature the DNA, with 35 cycles at 94 °C for 45 s, 50 °C for 60 s, and 72 °C for 90 s, with a final extension of 10 min at 72 °C to ensure complete amplification. Amplicons were quantified using PicoGreen (Invitrogen, Grand Island, NY, USA) assays and a plate reader, followed by clean-up using UltraClean® PCR Clean-Up Kit (Mo Bio Laboratories) and then quantification using Qubit readings (Invitrogen). The 16S rRNA gene samples were sequenced on an Illumina MiSeq platform (2 × 150 paired-end sequences) at Argonne National Laboratory core sequencing facility according to Earth Microbiome Project (EMP) standard protocols (Thompson et al. 2017).

### 16S rRNA gene pyrosequencing data preprocessing and analysis

Raw sequences were pre-processed, quality filtered and analyzed using the next-generation microbiome bioinformatics platform (QIIME2 v2021.4 pipeline) according to the developer’s suggestion (Hall and Beiko 2018). We used the DADA2 algorithm (Callahan et al. 2016), a software package wrapped in QIIME2, to identify amplicon sequence variants (ASVs) and perform quality control, such as filtering low-quality sequences, truncating to 120 base pair length, identifying and removing of chimera sequences, and merging paired-end reads to yield an ASV feature table (ASV table). Alpha and beta diversity analyses were performed in R using the *phyloseq* package (McMurdie and Holmes 2013). Alpha diversity was calculated by Shannon’s diversity index (Peet 1974a, Peet 1974b). Principal coordinate analysis (PCoA) was performed based on weighted and unweighted UniFrac distances, a method for computing differences between microbial communities based on phylogenetic information (Lozupone and Knight 2005). Weighted UniFranc considered both ASV presence and absence and abundance distances, and unweighted UniFrac only considered ASV presence. Permutational multivariate analysis of variance (PERMANOVA, R function adonis [vegan, 999 permutations]) was used to analyze paired statistical differences in beta diversity (Anderson 2001). Benjamini–Hochberg false discovery rate (FDR) correction was used to correct for multiple hypothesis testing (Storey 2002). For taxonomic comparisons, relative abundances based on all obtained reads were used. We used the QIIME2 plugin “q2-feature-classifier” and the Naïve Bayes classifier that was trained on the metagenome annotation package Greengenes13.8 99% operational taxonomic units (OTUs) full-length sequences to obtain the taxonomy for each ESV(DeSantis et al. 2006). Significantly different ASVs were determined using the analysis platform of Linear discriminant analysis effect size (LefSe) (Segata et al. 2011).

### Random Forest Model and Testing

16S metagenomic sequences from stool were analyzed on the QIIME2 platform (v2021.4). Briefly, samples were quality filtered and trimmed for PHRED>25. Samples were then denoised using the dada2 function, and the resulting feature counts table was rarefied to 4900 reads per sample. Taxonomy of ASVs was assigned using a classifier trained to the 515-806 region of the 16S rRNA gene acquired from the SILVA database (v138-99%). Feature selection and model generation were performed using the randomForest *(rf)* package in R. The ASV feature counts table for a subset of samples (20%) from weeks 4 and 8 was used as the initial training set and compared to the corresponding ALT levels for each sample at 24 weeks (24wkALT∼.) and a random forest model was generated using *ntree*=1000. Initial feature selection included diet as a categorical feature to compare the relative importance of a driving feature to the importance of ASVs but was removed from subsequent model testing (80%). Feature importance was calculated as a mean decrease in Gini scores and as a function of node purity. Additionally, features from the model were removed if they did not score above the importance score calculated after randomization of the data using the Boruta implementation of randomForest in R. The resulting model was then used in the R stats *predict* function for the remaining dataset of ASV counts from 4 and 8 weeks to generate predicted ALT scores based on the ASV abundances. Predicted ALT scores from the random forest model were compared to the actual ALT score through a linear regression for each treatment group and the entire dataset using the R stats ‘lm’ function. We used the python-based visual programming package orange v3 with the function ‘ROC analysis’ to evaluate our models by creating receiver operating characteristic (ROC) curves and calculating the area under the ROC curve (AURC). Model specificity and sensitivity were tested by converting the predicted and actual ALT values into binned values of LOW, MEDIUM, and HIGH before evaluation using the ‘*test and score’* module in *Orangev3*.*27*. We used the python package ‘poorly’, orange v3, and R ‘tidyverse’ package, including ggplot2, to visualize boxplots, scatterplots, and ROC curves.

### Construction of co-occurrence network development, calculation of centrality, and cluster analysis

We used Spearman to identify correlations of the abundance among bacterial ASVs. A correlation between two ASVs was considered statistically robust if the Spearman’s correlation coefficient (*ρ*) was >0.6 or <-0.6 with FDR *P*-value <0.05, resulting in robust pairwise correlations forming co-occurrence networks. The vertices in these networks represent ASVs and the edges that connect these vertices represent correlations between ASVs. We also relied on the R package “igraph” to analyze several topological metrics for each vertex of the co-occurrence networks. This feature set included betweenness centrality (the number of shortest paths going through a vertex), closeness centrality (the number of steps required to access all other vertices from a given vertex), and degree (the number of adjacent edges). Changes in these indices explain robustness of the network, the density of its links, or the vertices that tend to co-occur more frequently. An error and attack tolerance test was carried out by removing vertices of each network and measuring their respective loss of connectivity (Albert, Jeong and Barabási 2000).

### Metagenomic shotgun and data preprocessing and analysis

The DNA from stool samples was sent to Marine Biological Laboratory (US) for paired-end shotgun sequencing with up to 16 samples pooled in one lane using the Illumina NextSeq platform and generating 2 × 150 nt reads. Raw sequences obtained from metagenomic samples were run through Trimmomatic 0.39 (Bolger, Lohse and Usadel 2014) to remove low-quality base pairs, sequencing adapters, and reads shorter than 100 bp. To remove sequences of mouse origin, all reads were mapped to the mouse reference genome mm10 using Bowtie2 v.2.2.3 (Langmead and Salzberg 2012), and only the unmapped reads were used for further analysis. We retained only reads of at least 100 bp after trimming. After trimming, all libraries were interleaved for downstream analyses.

### Taxonomical analysis

For microbial profiling, the high-quality non-mouse reads were mapped against a set of clade-specific markers (spanning bacteria and archaea at the species level or higher) using the Metagenomic Phylogenetic Analysis (MetaPhlAn) tool v.3.0 and the marker database using default settings (Segata et al. 2012). To provide the relative abundance of each taxonomic unit, Bowtie2 v.2.2.3 was used for alignment (option “very-sensitive”), followed by normalization of the total number of reads in each clade by the nucleotide length of its marker.

### Functional profiling

To describe metabolic potential of the identified microbes, HMP (Human Microbiome Project) Unified Metabolic Analysis Network (HUMAnN3 v.3.0) was used with default settings (Abubucker et al. 2012). HUMAnN investigates presence/absence and relative abundances of gene families and pathways in each sample to provide a functional interpretation of the metagenomic sequences. HUMAnN was run for each sample separately using the taxonomic profiles from MetaPhlAn. For nucleotide-level searches, Bowtie2 v.2.2.3 was used to map reads to the functionally annotated pangenome database ChocoPhlAn. All unmapped reads were used for translated searches against the universal protein reference database UniRef50 (Wu et al. 2006) using Diamond v. 0.9.36 (Buchfink, Xie and Huson 2015). The resulting organism-specific gene hits were assigned to pathways through alignment to Kyoto Encyclopedia of Genes and Genomes (KEGG) pathways using MinPath v.1.2 (Ye and Doak 2009), finally providing a set of pathways and their relative abundances. To identify differences among diet treatments, LEfSe was applied to all pathways using default settings. A difference was statistically significant if p < 0.05 (Kruskal-Wallis test) and LDA score ≥ 2.

### Liver RNA sequencing and data preprocessing, processing, normalization, and differential expression analysis

The mouse liver samples were collected from GF and SPF mice after 8 and 24 weeks of dietary exposure. Three to six replicates were taken for each distinct condition. The RNA samples were sequenced on an Illumina NovaSEQ6000 platform via the genomics facility at the University of Chicago. The samples were sequenced by RNA-seq technology to produce raw reads in fastq format. The raw fastq files generated from the RNA-seq analysis were examined by the fastqc to evaluate the quality. The STAR (Dobin et al. 2013) (version 2.4.2a, Stanford University, CA) aligner was used to map the raw reads to the reference mouse genome (GRCm38 https://www.ncbi.nlm.nih.gov/grc). The STAR default parameter for the maximum mismatches is 10, which is optimized based on mammalian genomes and recent RNA-seq data. The genetic features from Gencode (Frankish et al. 2019) vM23 were extracted from the resulting bam file produced by STAR. The raw gene expression count matrix was generated by featureCounts (Chen, Lun and Smyth 2016) (version subread-1.4.6-p1). We used the EdgeR (Robinson, McCarthy and Smyth 2010) package (version 3.22.5) to filter low-expressed genes where genes expressed below ten counts-per-million (CPM) in 4 samples were removed. The Limma-Voom (Ritchie et al. 2015) pipeline (version 3.38.3) was applied to normalize the filtered count matrix and perform differential gene expression analysis for pairwise comparisons. In each comparison, we investigated the change of one experimental condition at a time, such as microbial status (GF vs SPF), time (8w vs 24w), or diet (FFVHC vs HF). We further examined the impact on gene expression profiles by diet changes and longitudinal variations. Differentially expressed genes (DEGs) were identified using FDR adjusted P-value cutoff of 0.05. Enrichment analysis on significant DEGs with mouse-specific GO terms and KEGG pathways was conducted through Lynx (Sulakhe et al. 2014) enrichment. Significantly enriched categories were selected using FDR adjusted P-value < 0.05 supported by Lynx API (Application Programming Interface). We showed the top 10 enriched categories in the bar plot sorted by the DEG hit ratio from the background mouse genome. Enriched GO terms and KEGG pathways were grouped into major categories based on immune and metabolism-related functions. The presence of dysregulated DEGs in the grouped GO terms or pathways is shown in the heatmap plot where up-and downregulated genes are distinguished by red and blue colors, respectively.

### Statistical Analysis

Statistical analysis was performed using GraphPad PRISM statistical software package (Mary L. Swift 1997). Data are presented as means +/- standard error of the mean (SEM), with statistical significance set at p< 0.05, 0.01, or 0.001. Statistical significance was determined via the nonparametric Kruskal-Wallis test followed by Turkey’s multiple comparisons, where p<0.05 was considered significant.

## Supplemental Figure Legends

**Figure S1.**
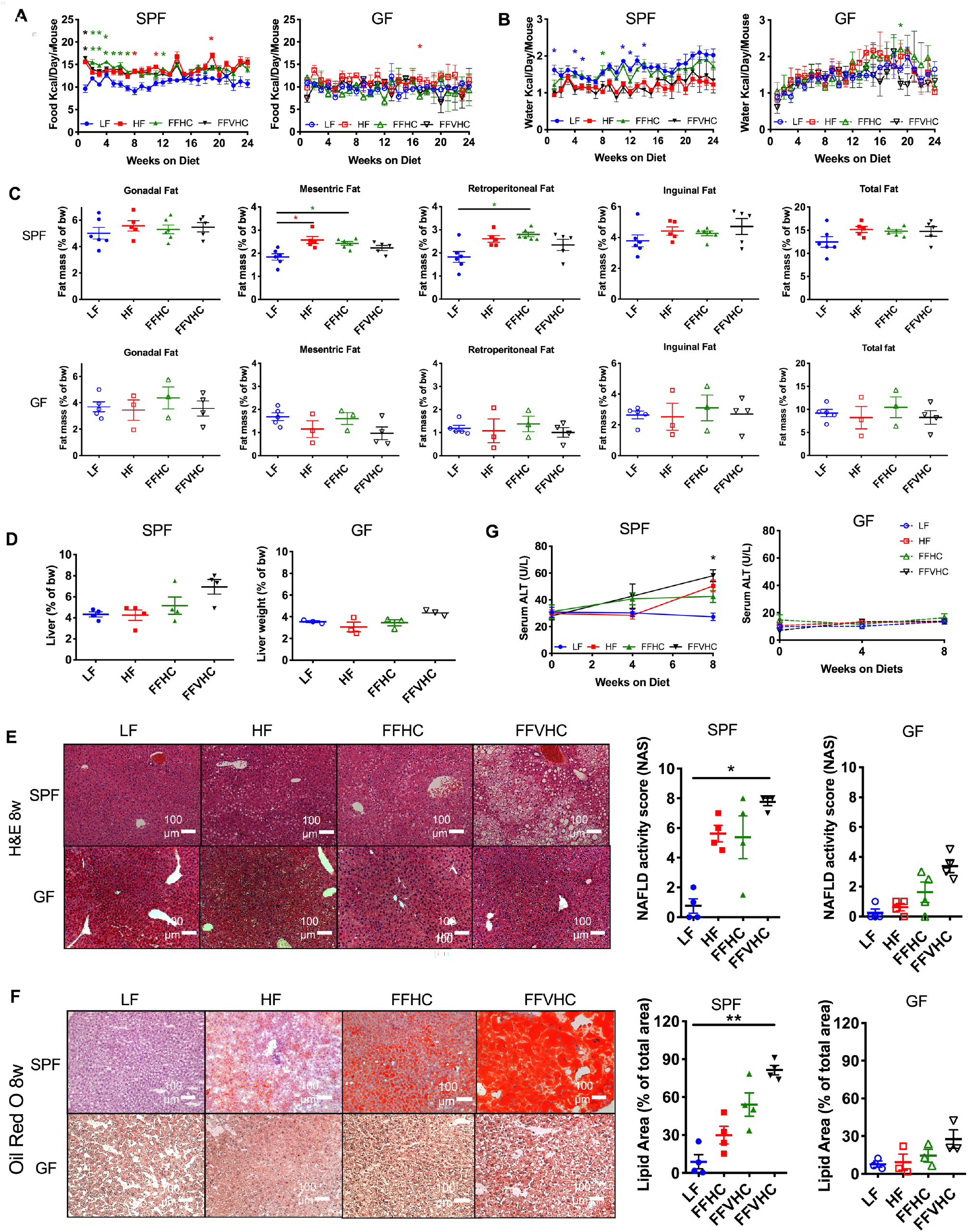
Dietary cholesterol interacts with gut microbes to differentially impact NAFLD/NASH development with corresponding features of disease occurring early and persisting late in disease in SPF and GF mice. (**A**) Food intake and (**B**) water intake throughout the 24 weeks. (**C**) Adipose tissue weight and (**D**) liver weight expressed as % of body weight in SPF and GF mice at 24 weeks. (**E**) Formalin-fixed liver hematoxylin and eosin stain (H&E, left) and corresponding graph representing NAFLD activity score (NAS, right) in SPF and GF mice at 8 weeks. (**F**) Oil Red-O staining (left) and corresponding graph representing lipid area quantification in SPF and GF mice at 8 weeks. Representative images are shown for livers from each group. All microscopy images were at 100x magnification, scale bars = 100 mm. (**G**) Alanine aminotransferase (ALT) levels in peripheral plasma of SPF and GF mice. n=4-6 per group. Data represent mean ± S.E.M. *P < 0.05, **P < 0.01, ***P < 0.001 via nonparametric Kruskal-Wallis test followed by Tukey’s multiple comparisons between means of all groups. Star color represents the treatment group exhibiting significant differences.

**Figure S2.**
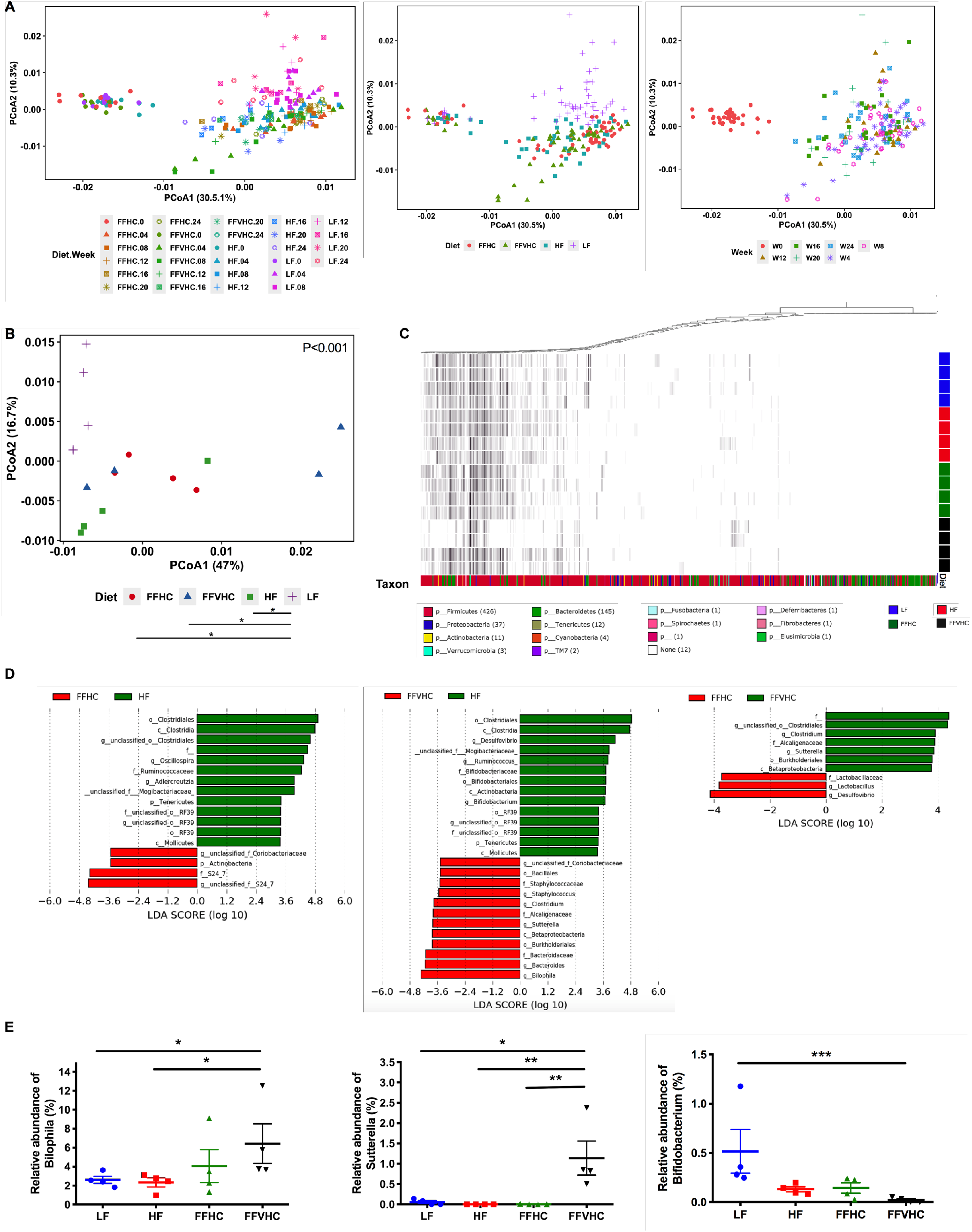
Fast food diets promote rapid gut dysbiosis, which precedes the onset of NAFLD. Beta diversity Principal Coordinate Analysis (PCoA) plots for Weighted Unifrac distances in (**A**) feces over time and (**B**) in cecal contents at week 8 (Axis 1 = principal coordinate (PC) 1, Axis 2 = PC 2). Small dots represent individual samples. (**C**) Heat map indicating relative abundances of Amplicon Sequence Variants (ASVs) in cecal bacteria composition induced by diet (week 8). Each column in the dendrogram represents an individual ASV. Each row represents an individual sample, organized by dietary treatment to the right of the heat map. Phyla-level taxonomic assignments are indicated by the colored bars at the bottom of the heat map. (**D**) Linear discriminant analysis effect size (LEfSe) analysis (LDA score [log10] bar graphs of differentially abundant taxa from the cecum of HF vs. FFHC, HF vs. FFVHC, and FFHC vs. FFVHC mice at week 8. Statistically significant taxa are presented (p<0.05) with an LDA score greater than ±3. Prefixes represent taxonomic rank, i.e., phylum (p), class (c), etc. (**E**) Relative abundance of significantly changed genera induced by FF diets identified by LEfSe with LDA score greater than ±3.0. n=4-6 per group. Data represents mean ± S.E.M. *P < 0.05, **P < 0.01, ***P < 0.001 via nonparametric Kruskal-Wallis test followed by Turkey’s multiple comparisons across all groups.

**Figure S3.**
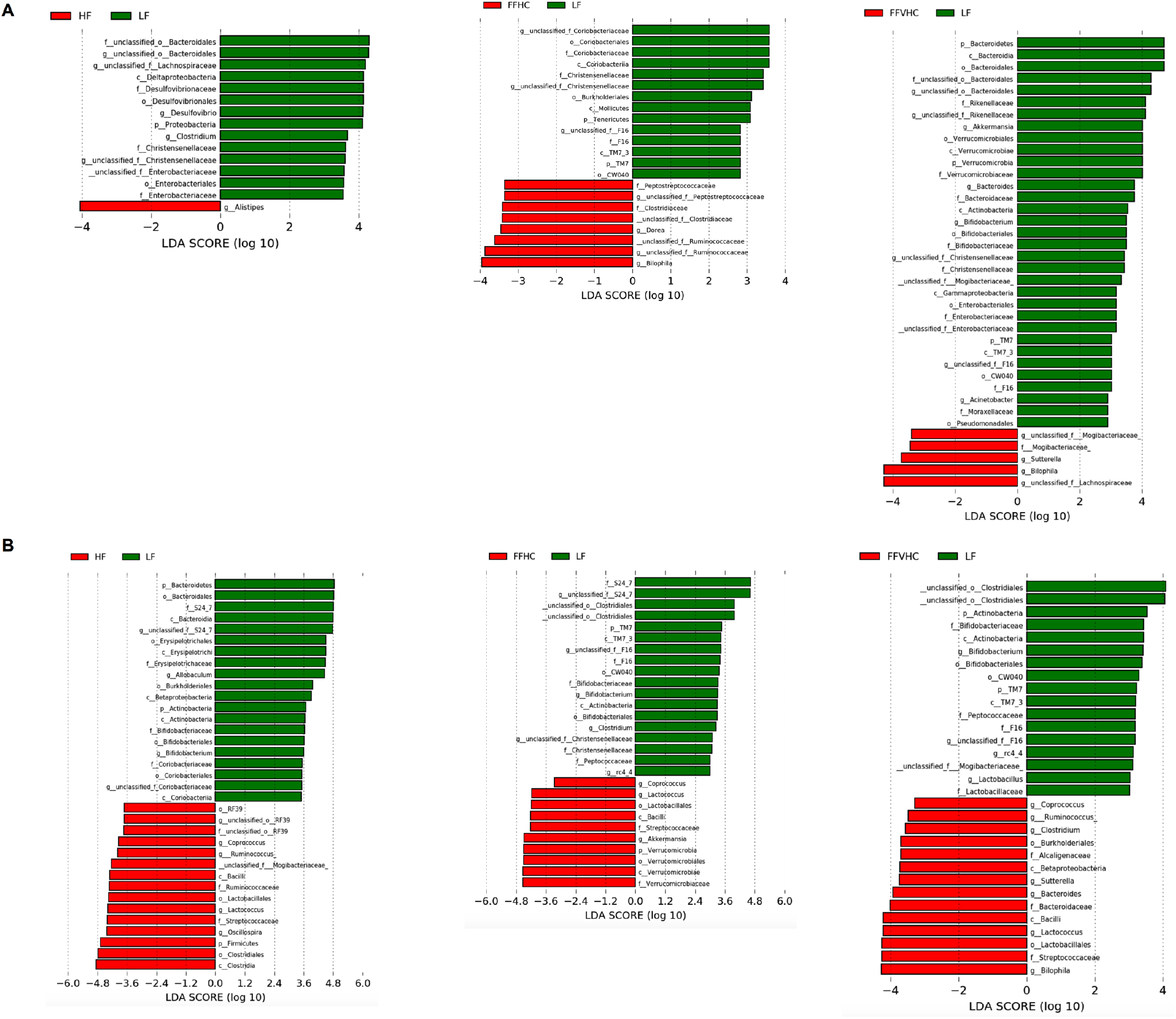
Fast food diets drive gut dysbiosis in the cecum that is accompanied by onset of NAFLD and NASH after 8 weeks of feeding. Linear discriminant analysis effect size (LEfSe) analysis (LDA score [log10] bar graphs of differentially abundant taxa from cecum of LF vs. HF, LF vs. FFHC, and LF vs. FFVHC at 24 (**A**) and 8 weeks (**B**). Statistically significant taxa are presented (p<0.05) with an LDA score greater than ±2. Prefixes represent taxonomic rank, i.e., phylum (p), class (c), etc. (**E**) Relative abundance of significantly changed genera induced by FF diets identified by LEfSe with LDA score greater than ±3.0.

**Figure S4.**
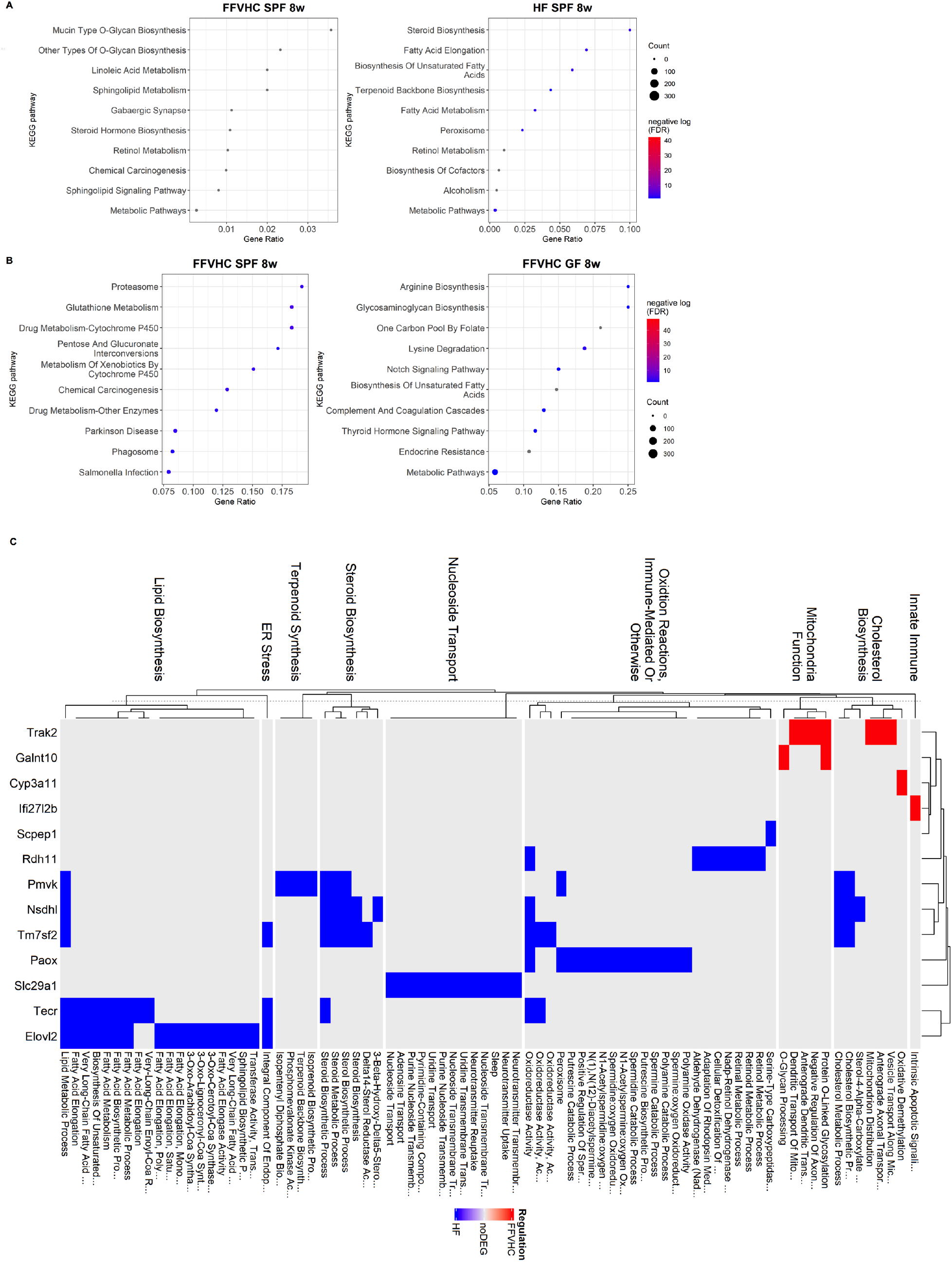
Dietary cholesterol level interacts with gut microbes to differentially impact systemic and local inflammation that begins early and persists late in disease in SPF and GF mice. (**A**) The top 10 enriched categories in the bar plot are sorted by DEG hit ratio from the background mouse genome after 8 weeks of feeding between SPF FFVHC-vs. SPF HF-fed mice **(A)** and between SPF FFVHC-vs. GF FFVHC-fed mice (**B**). (**C**) Enriched GO terms and KEGG pathways were grouped into major categories based on immune and metabolism-related functions, as shown in the heatmap plot. Up-and-down-regulated DEGs in the grouped GO terms or pathways are distinguished in red and blue colors, respectively. n=4-6 per group. Data are presented as mean± S.E.M. *P <0.05 via nonparametric Kruskal-Wallis test, followed by Turkey’s multiple comparisons between the means of all the groups. Star color represents treatment group exhibiting significant differences.

**Figure S5.**
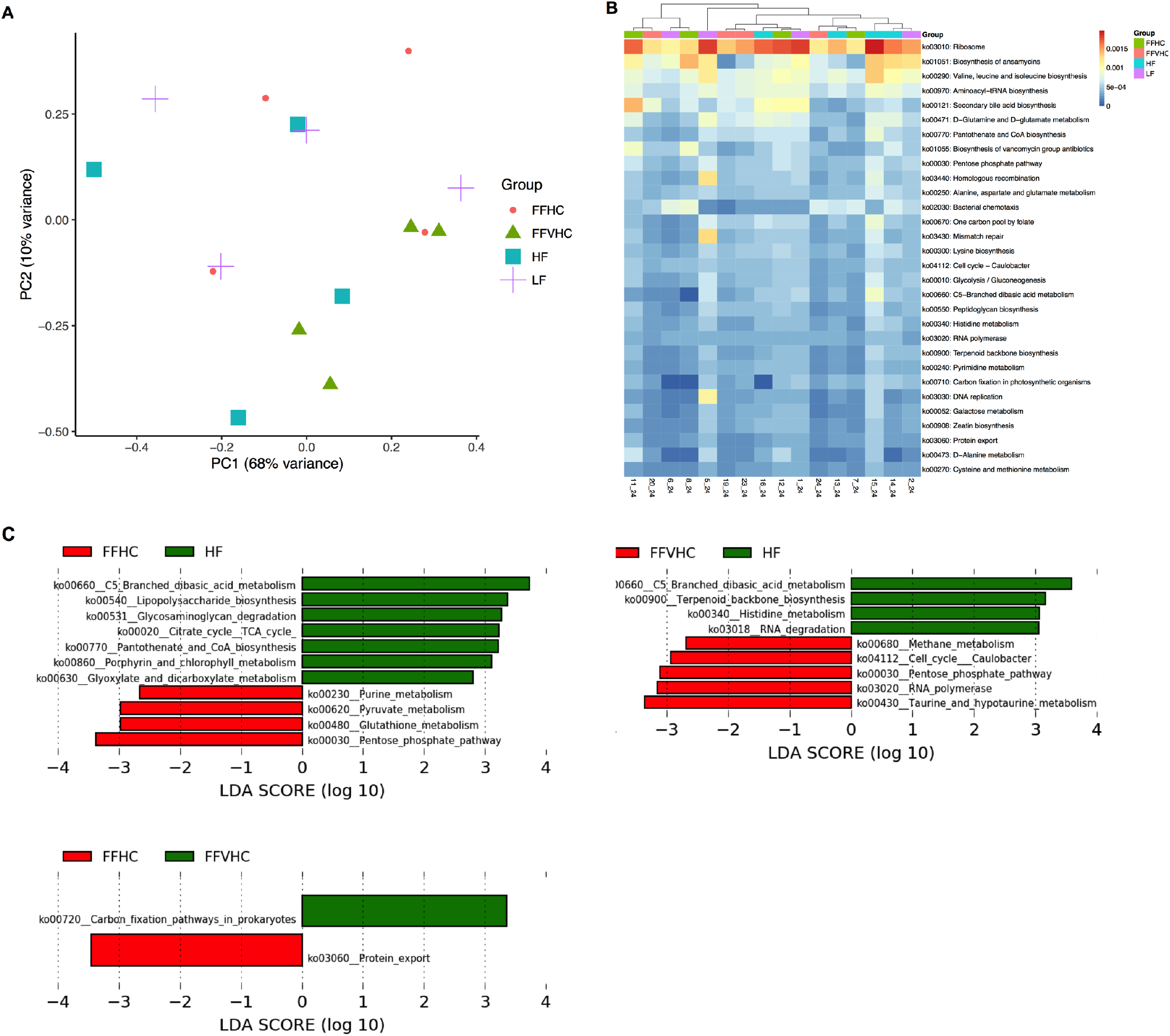
Differentially abundant pathways in fecal gut microbiota after 24 weeks and 8 weeks of feeding. (**A**) PCA plot for metabolic pathways in the gut microbiota induced by diets. **(B)** Heat map indicating relative abundances of metabolic functions in fecal bacteria induced by diets. Each column in the dendrogram represents an individual sample. Each row represents an individual functional pathway. Sample group assignments are indicated by the colored bars at the top of the heat map. (**C**) Linear discriminant analysis effect size (LEfSe) (LDA score [log10] bar graphs of differentially abundant metabolic functions from feces of HF vs. FFHC, HF vs. FFVHC, FFHC vs. FFVHC, LF vs. FFHC, LF vs. FFVHC, LF vs. HF mice at 24 weeks. Statistically significant taxa are presented (p<0.05) with an LDA score greater than ±3.

**Figure S6.**
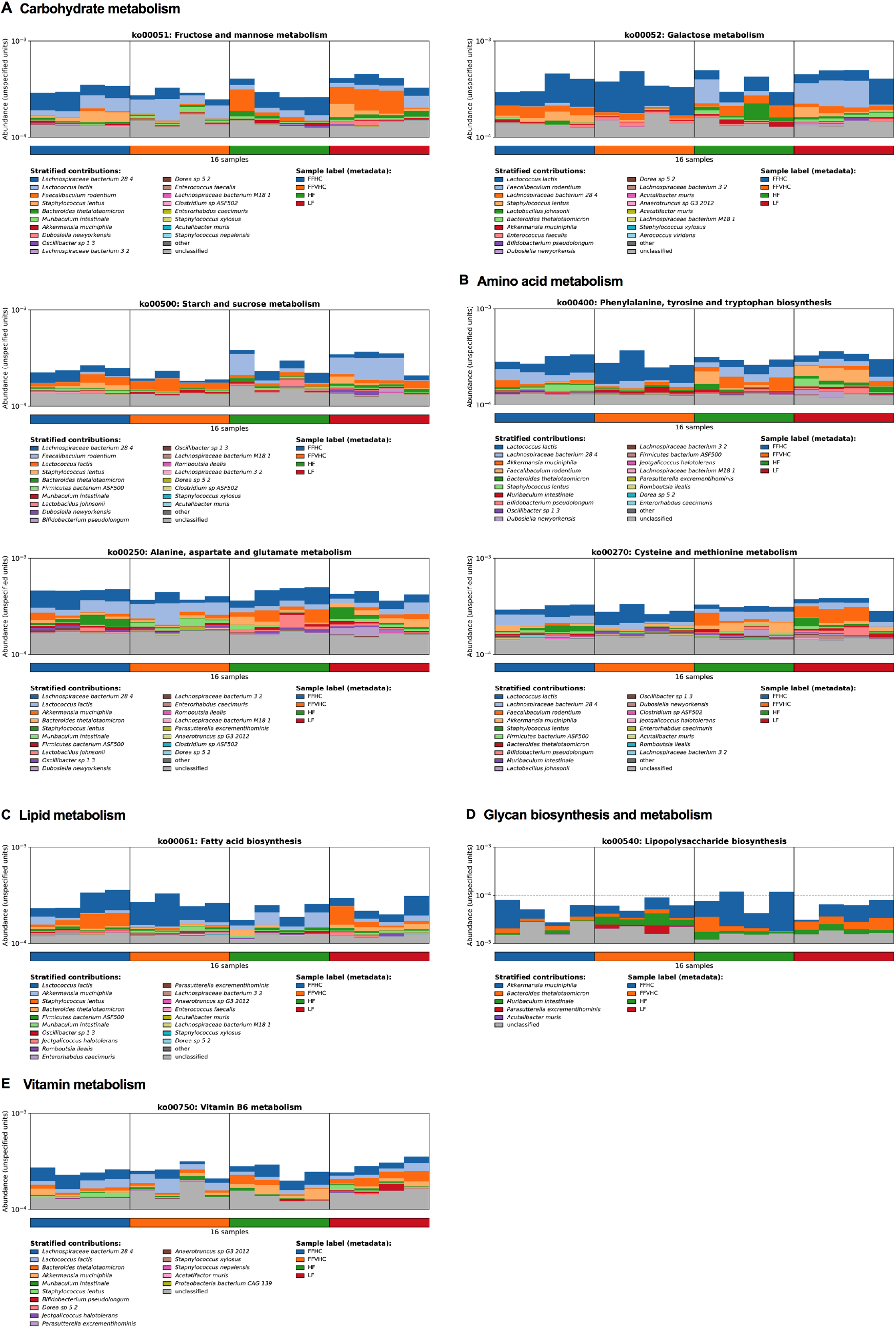
Differential metabolism in fecal gut microbiota after 24 and 8 weeks of feeding. (**A-E**) Microbiome metabolic features in each sample are separated based on MetaPhlAn taxonomic profiles for (**A**) Carbohydrate metabolism, (**B**) Amino acid metabolism, (**C**) Lipid metabolism, (**D**) Glycan biosynthesis and metabolism, and (**E**) Metabolism of vitamins.

